# Epoxy fatty acid dysregulation and neuroinflammation in Alzheimer’s disease is resolved by a soluble epoxide hydrolase inhibitor

**DOI:** 10.1101/2020.06.30.180984

**Authors:** Anamitra Ghosh, Michele E. Comerota, Debin Wan, Fading Chen, Nicholas E. Propson, Sung Hee Hwang, Bruce D. Hammock, Hui Zheng

## Abstract

Neuroinflammation has been increasingly recognized to play critical roles in Alzheimer’s disease (AD). The epoxy fatty acids (EpFAs) are derivatives of the arachidonic acid metabolism with anti-inflammatory activities. However, their efficacy is limited due to the rapid hydrolysis by the soluble epoxide hydrolase (sEH). We found that sEH is predominantly expressed in astrocytes where its levels are significantly elevated in postmortem human AD brains and in β-amyloid mouse models, and the latter is correlated with drastic reductions of brain EpFA levels. Using a highly potent and specific small molecule sEH inhibitor, 1-trifluoromethoxyphenyl-3-(1-propionylpiperidin-4-yl) urea (TPPU), we report here that TPPU treatment potently protected against LPS-induced inflammation in vitro and in vivo. Long-term administration of TPPU to the 5xFAD mouse model via drinking water reversed microglia and astrocyte reactivity and immune pathway dysregulation, and this is associated with reduced β–amyloid pathology and improved synaptic integrity and cognitive function. Importantly, TPPU treatment reinstated and positively correlated EpFA levels in the 5xFAD mouse brain, demonstrating its brain penetration and target engagement. These findings support TPPU as a novel therapeutic target for the treatment of AD and related disorders.

**One Sentence Summary:** We show that soluble epoxide hydrolase is upregulated in AD patients and mouse models, and that inhibition of this lipid metabolic pathway using an orally bioavailable small molecule inhibitor is effective in restoring brain epoxy fatty acids, ameliorating AD neuropathology and improving synaptic and cognitive function.

## Introduction

Alzheimer’s disease (AD) is the most common form of age-associated neurodegenerative disorder and is an unmet medical need. AD is defined by the deposition of extracellular senile plaques composed of amyloid beta (Aβ) aggregates and the formation of intracellular neurofibrillary tangles containing abnormal hyperphosphorylated tau protein (*1*). Overwhelming evidence support the notion that the accumulation of Aβ initiates a series of downstream events leading to cognitive impairment and neurodegeneration (*2*, *3*). Hence, the majority of AD clinical trials have focused on reducing Aβ load. Unfortunately, these trials have been unsuccessful so far (*2*, *4*–*6*). Thus, there is an urgent need to pursue other disease modifying therapies.

Besides the pathological hallmarks, AD is associated with prominent neuroinflammation (*7*, *8*). Prolonged activation of glial cells, microglia and astrocytes in particular, and the release of proinflammatory cytokines, chemokines, and reactive oxygen and nitrogen species, create a neurotoxic environment which could exacerbate the progression of AD (*9*–*11*). Recent genome wide association studies have identified multiple immune related gene variants as risk factors for late-onset AD, supporting a major contributing role of innate immunity and neuroinflammation in AD (*12*–*15*). However, while epidemiological studies indicated positive effects for nonsteroidal anti-inflammatory drugs (NSAIDs) in AD development, randomized clinical trials failed to demonstrate clinical efficacy (*16*, *17*).

Arachidonic acid (ARA) is an omega-6 unsaturated fatty acid present in the plasma membrane where it is bound to phospholipids (*18*). It can be released from the membrane by phospholipase A2 (PLA_2_) and further metabolized by enzymes in three major pathways: cyclooxygenases (COXs), lipoxygenases (LOXs), and cytochrome P450 enzymes (CYPs), which produce prostaglandins, leukotrienes, and various eicosanoids including epoxyeicosatrienoic acids (EETs) (*19*, *20*). Among these the COX and LOX pathways have been extensively studied and successfully targeted therapeutically (*19*). Of note, most of the NSAIDs are COX-1 and/or COX-2 inhibitors. In contrast, much less is known about the therapeutic potential of the CYP pathway.

Distinct from the well-established proinflammatory role of the prostaglandins, EETs and other epoxy fatty acids (EpFAs) have been proposed to possess anti-inflammatory properties (*21*, *22*). However, they are broken down rapidly into their corresponding diols by the soluble epoxide hydrolase (sEH). Genetic deletion of *Ephx2* (gene encoding sEH) or pharmacological inhibition of sEH conferred beneficial effects in several disease models, including depression, Parkinson’s disease (*23*–*26*), and most recently, APP/PS1 transgenic mouse model of AD (*27*). However, these studies are restricted to acute model systems or germline deletions. It is not clear whether the sEH pathway can be therapeutically targeted under chronic conditions.

Here we present evidence that sEH is aberrantly elevated in the brain of AD individuals and Aβ mouse models, the latter is correlated with significant reduction of EpFA levels. Using a highly selective and potent sEH inhibitor, 1-trifluoromethoxyphenyl-3-(1-propionylpiperidin-4-yl) urea (TPPU) (*23*, *26*), we show that long-term administration of TPPU to the 5xFAD mouse model restored the EpFA levels and reversed microglia and astrocyte reactivity and their associated molecular signatures. These are accompanied by attenuated β–amyloid pathology and improved synaptic integrity and cognitive function.

## Results

### Elevated sEH and diminished EpFA levels associated with AD

We first evaluated the expression of genes involved in arachidonic acid (ARA) metabolism in postmortem AD brain samples and their age-matched healthy controls (Fig. 1, A-B). Quantitative real-time PCR (qPCR) analysis showed that the expression of *PLA2G2A* which catalyzes the release of ARA from the membrane phospholipids, but not fatty acid amide hydrolase (*FAAH*) which facilitates ARA production from endocannabinoids and degrades bioactive fatty acid amides, was significantly elevated, indicating that PLA2-mediated release of ARA is a major source for the upregulated ARA in AD. Both the *COX-2* and *CYP4F8*, which produce prostaglandin (PG) and its metabolite PGE_2_, respectively, were significantly increased, suggesting overall activation of the cyclooxygenase metabolism. Interestingly, examination of the CYP monooxygenase pathway revealed that, while no differences in several of the CYPs, including *CYP2J2*, *CYP2C8* and *CYP2C19*, were detected, expression of *EPHX2* was significantly higher in AD samples compared to the controls (Fig. 1B). These results suggest aberrant regulation of the ARA metabolism in AD brains.

**Fig. 1.**
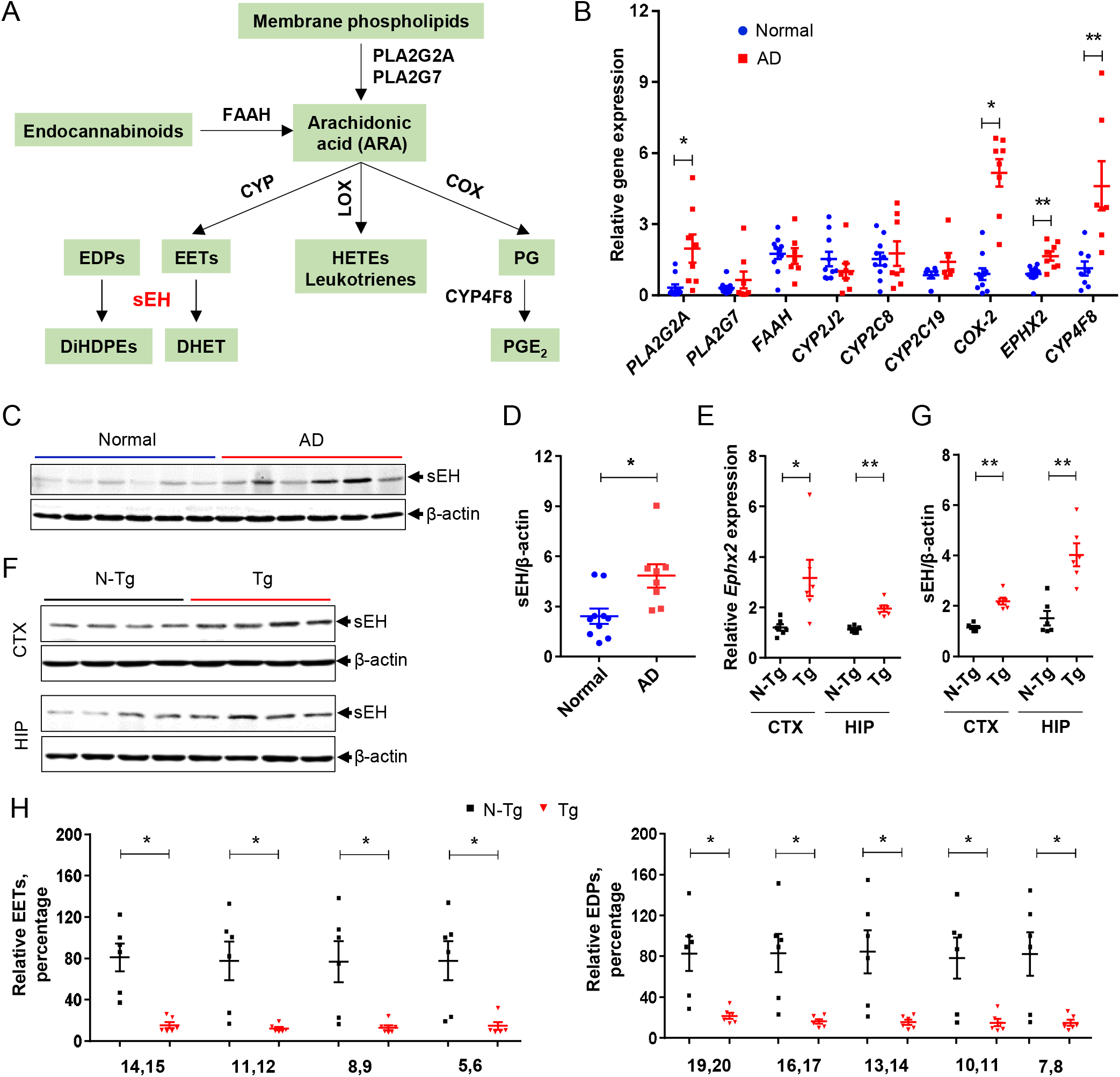
sEH and arachidonic acid (ARA) metabolism dysregulation in AD. (A) Schematic diagram of ARA metabolism pathway. (B) qPCR analysis of mRNA expression of ARA pathway genes in the brains of AD patients (n=8) and age-matched non-demented controls (n=10). (C) Representative Western blot illustrating the expression of sEH in human brains. β-actin was used as a loading control. (D) Quantification of sEH/β-actin ratio. (E) qPCR analysis of *Ephx2* in cortex (CTX) and in hippocampus (HIP) of littermate non-transgenic (N-Tg) and 5xFAD transgenic (Tg) mice at 4.5 months of age. (F) Representative Western blot illustrating the levels in CTX and HIP of Tg mice and N-Tg controls. β-actin was used as a loading control. (G) Quantification of sEH/β-actin ratio in CTX and in HIP. (H) Quantification of relevant EET and EDP regioisomers (displayed as percentage to a reference N-Tg mouse) in N-Tg and Tg mouse brains. The numbers at x-axis denote carbon numbers where the double bonds were located in the corresponding polyunsaturated fatty acids. Data are means ± SEM of eight to ten human brains per group (B-D) or six to eight mice per group (E-H). **P < 0.01, *P < 0.05. Data were analyzed by unpaired Student’s *t*-test.

Consistent with the mRNA expression, Western blot analysis revealed a nearly two-fold increase of sEH protein in AD brains compared to controls (Fig. 1C and quantified in D). To substantiate these findings, we performed qPCR analysis of 5xFAD transgenic (Tg) mice, which showed increased *Ephx2* expression in both the cortex and hippocampus at 4.5 months of age compared to their littermate non-transgenic (N-Tg) controls (Fig. 1E). Western blotting validated increased sEH levels in these mice (Fig. 1F and quantified in G). Similar increases were also detected in an APP^NLGF^ knock-in mice (*28*) with physiological expression of APP when compared with wild-type controls (fig. S1, A-B). The results combined demonstrate prominent upregulation of sEH in the brains of AD patients and APP/Aβ mouse models.

Consistent with augmented sEH levels, LC-MS/MS–based lipidomic analysis of two major sEH eicosanoid substrates, EETs and epoxydocosapentaenoic acids (EDPs) showed dramatically reduced levels of multiple EET and EDP regioisomers in Tg brains in comparison to N-Tg controls (Fig. 1H). Similar to that of human samples, no alterations in the expression of CYP monooxygenases that produce EETs from the ARA, such as *Cyp2c39*, *Cyp2j5* and *Cyp2j9*, were observed in Tg mice (fig. S1C). Thus, the reduced EETs and EDPs are likely due to specific increases of sEH activity downstream of CYP. These results suggest that elevated sEH plays an active role in hydrolyzing EETs and EDPs, leading to their reduced levels in Tg, and possibly human AD brains.

### TPPU blocks astroglial sEH upregulation and LPS-induced inflammation

To assess the cell type expression of sEH, we used a flow cytometry-based <u>co</u>ncurrent <u>br</u>ain cell type acquisition (CoBrA) method to simultaneously isolate astrocytes, microglia and vascular endothelial cells (*29*). qPCR analysis of *Ephx2* revealed that sEH is highly expressed in sorted astrocytes where its levels are significantly elevated compared to N-Tg controls (Fig. 2A). Interestingly, although modest and low levels of *Ephx2* were detected in endothelial cells and microglia, respectively, there were no significant differences between Tg and N-Tg controls. This result was substantiated by co-immunofluorescence staining of sEH with astrocyte (GFAP) or microglia (Iba-1) markers in hippocampal sections of N-Tg and Tg brains (Fig. 2B). We observed elevated expression of sEH predominantly in GFAP-positive astrocytes in Tg samples, with negligible co-staining with Iba1, consistent with previous findings (*27*). Both the sEH fluorescence intensity and the number of sEH expressing cells were increased significantly in Tg mice compared to N-Tg controls (Fig. 2C). Similar astrocytic upregulation of sEH was also observed by acute LPS administration in vivo (fig.S2) and in primary astrocyte cultures (fig. S3, A-B), and this is associated with increased nitrite release as measured by the Griess assay (Fig. 2D) and the expression of proinflammatory cytokines (Fig. 2E).

**Fig. 2.**
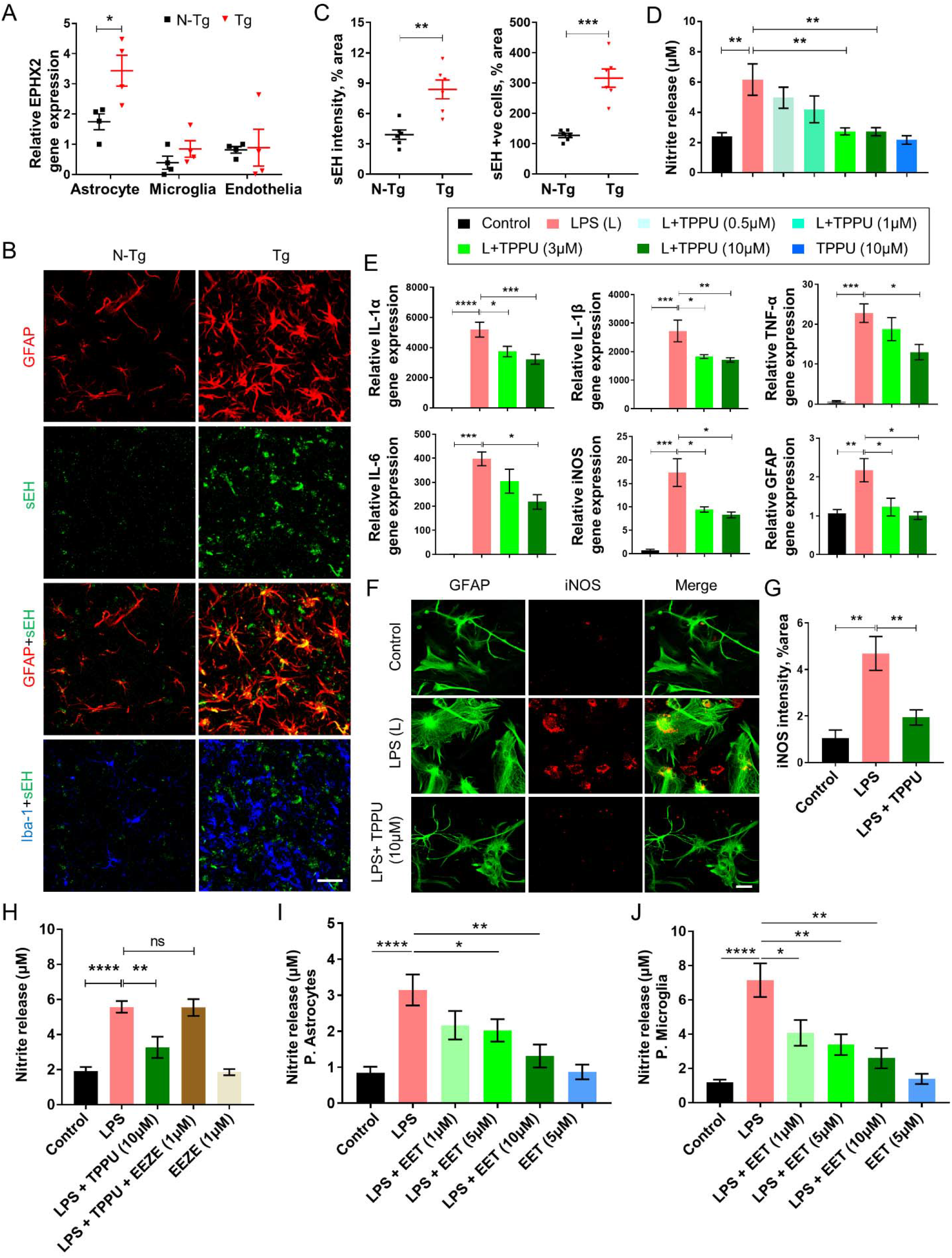
Astrocytic sEH upregulation in Tg mice and by LPS treatment. (A) qPCR analysis of mRNA expression of *Ephx2* in sorted astrocyte, microglia and endothelia of N-Tg and Tg mice. (B) Immunostaining of GFAP (red), sEH (green) and Iba-1 (blue) in the hippocampus of Tg mice at 4.5 months of age. Scale bar 50 μm. (C) Quantification of sEH intensity (left) and sEH+ve cells (right) from (B). (D-G) Analysis of conditioned media or cell lysates of primary astrocyte cultures pretreated with different doses of TPPU (ranging from 0.5 μM to 10 μM) for 30 minutes followed by LPS treatment (100 ng/ml) for 24 h. (D) Nitrite measurement from cultured media by Griess assay. (E) qPCR analysis of mRNA expression of *Il-1α*, *Il-1β*, *Tnf-α*, *Il-6*, *iNOS* and *Gfap*. (F) Immunocytochemistry of GFAP (green) and iNOS (red) in primary astrocytes. Scale bar, 100 μm. (G) Quantification of iNOS intensity in primary astrocytes. (H) Nitrite measurement of conditioned media from primary astrocytes pretreated with TPPU (10 μM) or pan-EET receptor antagonist 14,15-EEZE (1 μM) for 30 minutes followed by LPS treatment (100 ng/ml) for 24 hours. (i and j) Nitrite levels in conditioned media of primary astrocytes (I) or primary microglia (J) pretreated with different doses of 11,12-EET (ranging from 1 μM to 10 μM) for 30 minutes followed by LPS treatment (100 ng/ml) for 24 h. Data are means ± SEM of either four to six mice per group (A-C) or three independent experiments (n=3, D-J). ****P < 0.0001, ***P < 0.001, **P < 0.01, *P < 0.05, ns= not significant. Data were analyzed either by unpaired Student’s *t*-test (A-D) or by one-way ANOVA with Tukey’s multiple comparison test (D-J).

Using these parameters, we tested the effect of sEH inhibitor TPPU in LPS-treated mouse primary astrocyte cultures. Thirty minutes pre-treatment of TPPU dose-dependently reduced nitrite release (Fig. 2D), and the expression of proinflammatory molecules *Il-1α*, *Il-1β*, *Tnf-α*, *Il-6*, *iNOS* and *Gfap* (Fig. 2E) and *Ccl-2* and *Cxcl-1* (fig. S3C). The reduced expression of iNOS and GFAP were confirmed by immunostaining (Fig. 2, F-G). In agreement with the astroglial specific expression of sEH, TPPU failed to attenuate LPS-induced expression of pro-inflammatory molecules in primary microglia cultures (fig. S4).

### EETs are functional mediators of TPPU

Next, we wondered whether the anti-inflammatory effect of sEH inhibition was attributed to increased EETs. We pre-treated primary astrocytes with TPPU and/or a putative pan-EET receptor antagonist 14,15-EEZE 30 minutes prior to LPS treatment and measured the nitrite levels by Griess assay (*30*). We observed that the inhibitory effect of TPPU was completely abolished upon co-treatment with 14,15-EEZE (Fig. 2H), suggesting that EETs and possibly related EpFA are the functional lipid mediators of TPPU.

We then went on to test whether exogenous EET could directly mitigate LPS-induced inflammation. We found that 30-minute pre-treatment of 11,12-EET dose dependently attenuated the LPS-induced nitrite release in both cultured primary astrocytes (Fig. 2I) and microglia (Fig. 2J). EET treatment also suppressed the LPS-induced expression of pro-inflammatory molecules *Il-1α*, *Il-1β*, *Tnf-α*, *Il-6*, *iNOS* and *C3* in primary microglia cultures (fig. S5). The efficacy of EET in preventing LPS-induced cytokine expression was further validated using an ex vivo mouse hippocampal organotypic slice culture system (fig. S6). Collectively, these results support a pathway whereby TPPU inhibits astroglial sEH activity, and the resulting augmentation of EETs exert anti-inflammatory effect in both astrocytes and microglia through autocrine and paracrine activities.

### TPPU mitigates LPS-induced acute inflammation in vivo

Given the strong anti-inflammatory effect of TPPU in vitro, we thought to assess its in vivo efficacy. We first examined whether TPPU is a substrate for P-glycoprotein (P-gp), which mediates the ATP-dependent efflux of drugs or xenobiotics (*31*). Using the well-established human Caco-2 cells (*32*, *33*), we determined that the apparent permeability coefficient (*P_app_*) for TPPU from basolateral to apical (B to A) was 24.45 and from A to B was 18.03 (Table S1). Thus, the B to A/A to B efflux ratio was 1.36, which was decreased to 0.99 in the presence of verapamil (a P-gp inhibitor). Since a ratio of >2 is generally considered to involve P-gp-mediated efflux (*33*), TPPU is unlikely to be a good P-gp substrate.

Next, we investigated the effect of TPPU in LPS-induced inflammation by pre-treating the C57BL/6 mice with one dose of TPPU (3 mg/kg) via oral gavage for 24 hours, followed by co-treatment with LPS (3 mg/kg, i.p.) and TPPU (3 mg/kg, oral gavage), and the mice were euthanized after 18 hours (fig. S7A). Consistent with the in vitro and ex vivo studies, qPCR analysis of brain samples showed that LPS triggered the expression of proinflammatory molecules in both the cortex and hippocampus, and the vast majority of these were significantly reduced by TPPU (fig. S7B). Western blotting showed that levels of iNOS, COX-2, GFAP, Iba-1 and sEH were upregulated in LPS-treated mice but downregulated by TPPU in both the hippocampus and cortex (fig. S7, C-F). Immunofluorescence staining of Iba-1 and GFAP and co-staining with COX-2 and iNOS documented that TPPU mitigated LPS-induced microglia and astrocyte cell numbers and staining intensities, respectively, as well as COX-2 and iNOS expressions (fig. S8). Together, our results demonstrate that sEH blockade by TPPU prevents acute neuroinflammation in vitro and in vivo.

### TPPU enters the brain and engages its target under chronic treatment conditions

Given the heightened expression of sEH in AD human brains and mouse models and the strong acute anti-inflammatory effect of TPPU, we thought to test the long-term therapeutic effect of TPPU in 5xFAD Tg mice. We supplied either Vehicle (Veh) or TPPU to Tg mice and their N-Tg littermate controls via drinking water starting at 2 months of age and continuously for 2.5 or 4.5 months (fig. S9A). Measurement of average water consumption per week per mouse for 10 weeks found no significant differences between Tg Veh and Tg TPPU groups, demonstrating that TPPU did not affect fluid intake (fig. S9B). Measurement of TPPU in brain and plasma samples showed that, whereas TPPU is undetectable in Veh-treated controls, its levels can be clearly measured both in the brain (fig. S9C) and plasma (fig. S9D); the resulting brain to plasma ratio is 21.7% and 17.2% for N-Tg and Tg group, respectively, consistent with that of acute administration (*26*). These results establish that TPPU is able to gain access to the brain where its levels can be maintained under chronic treatment conditions.

To determine the target engagement of TPPU in the brain, we measured levels of EETs and EDPs, which were significantly reduced in Tg brains (Fig. 1H). We found that the EETs (Fig. 3A) and EDPs (Fig. 3B) were elevated in TPPU-treated Tg mice compared to the vehicle controls. The trending, but not statistically significant, increases of some of the regioisomers could be attributed to the differences in drug uptake or response among individual animals. Interestingly, plotting the co-expression relationship between EpFA regioisomers and TPPU observed prominent positive correlations between these two factors in the brain (Fig. 3, C-D), strengthening the notion that TPPU penetrates the brain and antagonizes its target sEH.

**Fig. 3.**
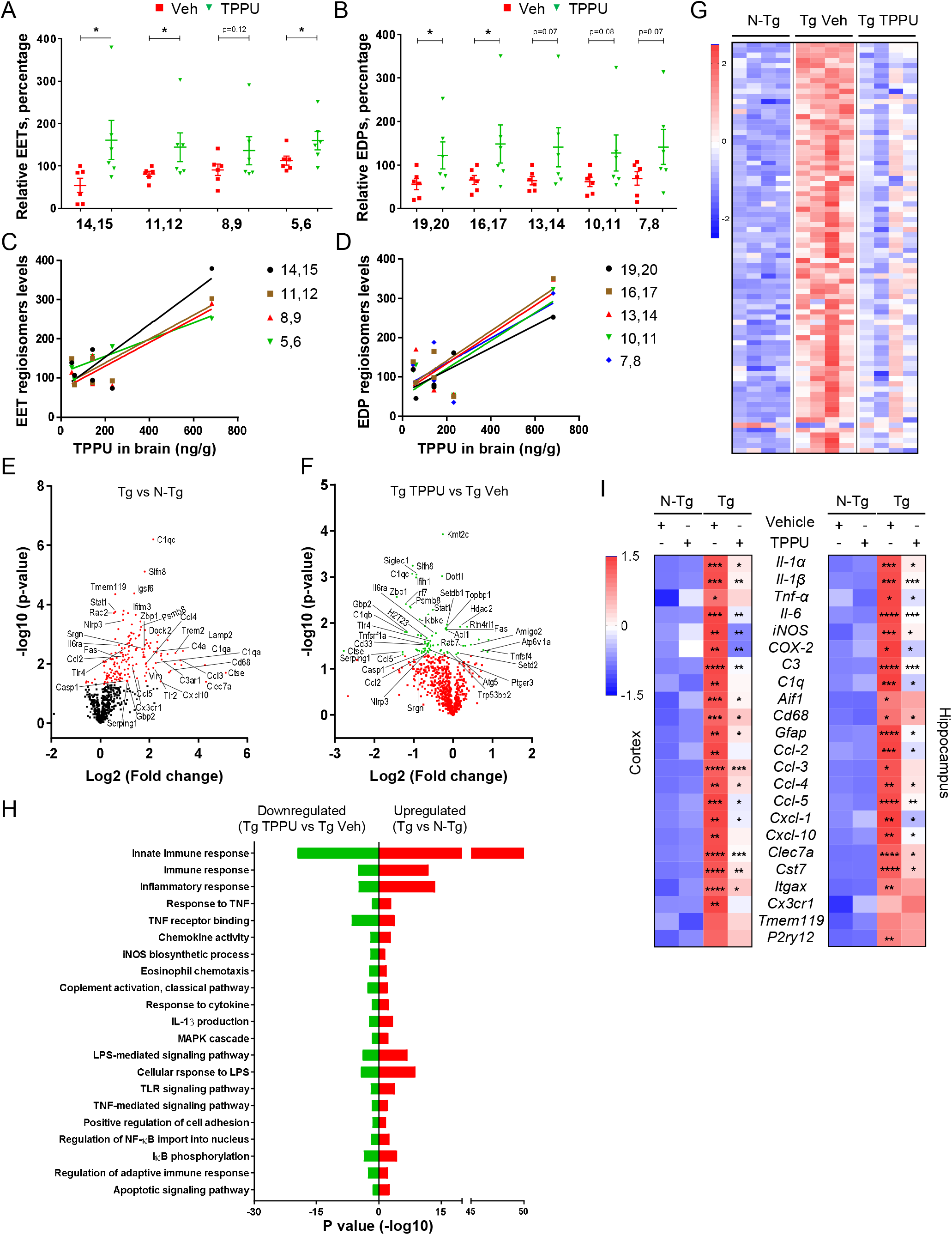
TPPU engages its target and attenuates inflammatory gene expression in Tg mice. (A, B) Quantification of relevant EETs (A) and EDPs (B) (displayed as percentage to a reference Tg Veh mouse) from vehicle (Veh) and TPPU-treated mouse brains. (C) Correlation of brain TPPU with different EET regioisomers (r=0.874, p<0.0238 for 14,15-EET; r=0.866, p<0.026 for 11,12-EET; r=0.02 for 8,9-EET; r=0.947, p<0.004 for 5,6-EET). (D) Correlation of brain TPPU with different EDP regioisomers (r=0.899, p<0.015 for 19,20-EDP; r=0.84, p<0.037 for 16,17-EDP; r=0.796, p<0.581 for 13,14-EDP; r=0.84, p<0.036 for 10,11-EDP; r=0.762, p<0.079 for 7,8-EDP). (E, F) Volcano plot of Nanostring neuroinflammation gene expression profiling showing differences in gene expression in the hippocampus stratified by (E) Tg vs N-Tg and (F) Tg TPPU vs Tg Veh. For each plot, significance is plotted against fold-change (log2 values). Red dots and green dots denote genes with adjusted significance of p<0.05. (G) Heat map of relative expression of inflammatory pathway genes in N-Tg, Tg Veh and Tg TPPU mice. (H) Gene ontology (GO) and pathway analysis of differentially expressed genes (DEGs). Significant GO terms (biological processes) associated with identified DEGs. The vertical axis represents the GO category, and the horizontal axis represents the P-value (–Log10) of the significant GO terms. Red bars represent the significantly upregulated inflammatory pathways in Tg mice vs N-Tg mice and green bars represent significantly downregulated inflammatory pathways in Tg TPPU mice vs Tg Veh mice. (I) Heat map showing qPCR analysis of mRNA expression in cortex (left) and in hippocampus (right). The asterisks in Tg: Vehicle (+) TPPU (−) column represent significant change vs N-Tg: Vehicle (+) TPPU (−). The asterisks in Tg: Vehicle (−) TPPU (+) represent significant change vs Tg: Vehicle (+) TPPU (−). Data are means ± SEM of either six to eight mice (A-D) or four mice (E-H) per group. Data were analyzed by Student’s *t*-test or by two-way ANOVA with Bonferroni’s multiple comparison test. For C and D, correlation coefficients (r) were computed using Pearson correlations and each dot represents individual mouse and linear regression line (solid).

### TPPU treatment reverses immune pathway dysregulation in Tg mice

Having established the pharmacodynamics of TPPU, we tested its effect in 5xFAD mice by employing molecular, biochemical, neuropathological and functional approaches as outlined (fig. S9A). To investigate the molecular mechanisms, we performed multiplex gene expression analysis using a Nanostring nCounter panel enriched for inflammatory genes. We quantified expression of 757 genes from the hippocampi of 4.5-month-old N-Tg and Tg mice treated with Veh or TPPU (n=4/group). Volcano plot representation of gene expression stratified by Veh-treated Tg vs N-Tg (Fig. 3E) demonstrated significant upregulation of 171 genes (red dots) in Tg mice. For each dot, significance is plotted against fold-change (log2 values). Treatment with TPPU for 2.5 months downregulated 73 inflammatory genes (green dots) compared to the Tg Veh group (Fig. 3F). Analysis of eigengene expression by heatmap in this inflammatory module for individual animals demonstrated that increased inflammatory gene expression in the Tg Veh was overall downregulated by TPPU (Fig. 3G). Next, we performed gene ontology (GO) enrichment analysis to gain further insights into the biological functions of differentially expressed genes (DEGs). Our results showed that 64 pathways were significantly enriched for the identified DEGs (P < 0.05). DEGs that were mainly enriched in immune-related processes, such as immune responses, inflammatory responses, chemokine responses, cytokine mediated signaling pathway, iNOS biosynthetic process, complement activation pathway, LPS-mediated signaling pathway, cell adhesion and chemotaxis pathway, NF-ĸB mediated signaling pathway and apoptotic signaling pathways (Fig. 3H, Tg vs N-Tg). All the inflammatory pathways upregulated in Tg Veh mice were significantly downregulated with TPPU treatment (Fig. 3H, Tg TPPU vs Tg Veh). The Nanostring results were further validated by qPCR analysis of selected proinflammatory molecules using both cortex and hippocampal tissues (Fig. 3I).

In line with the gene expression data, Western blot analysis documented elevated levels of inflammatory (iNOS and COX-2) and glial cell (GFAP and Iba-1) markers in Tg mice compared to the N-Tg controls both in hippocampus (Fig. 4, A-B) and cortex (fig. S9, E-F). 2.5 months of TPPU treatment resulted in significant downregulation of these proteins. These results were further validated by immunofluorescence staining of Iba-1 (Fig. 4C) and GFAP (Fig. 4D) and co-staining with COX-2 and iNOS, followed by quantification of their intensities and the number of Iba-1 and GFAP positive cells (Fig. 4E). Together, the findings strongly suggest that inhibition of sEH by TPPU is effective in reversing the dysregulated immune pathways and glia reactivity in Tg mice.

**Fig. 4.**
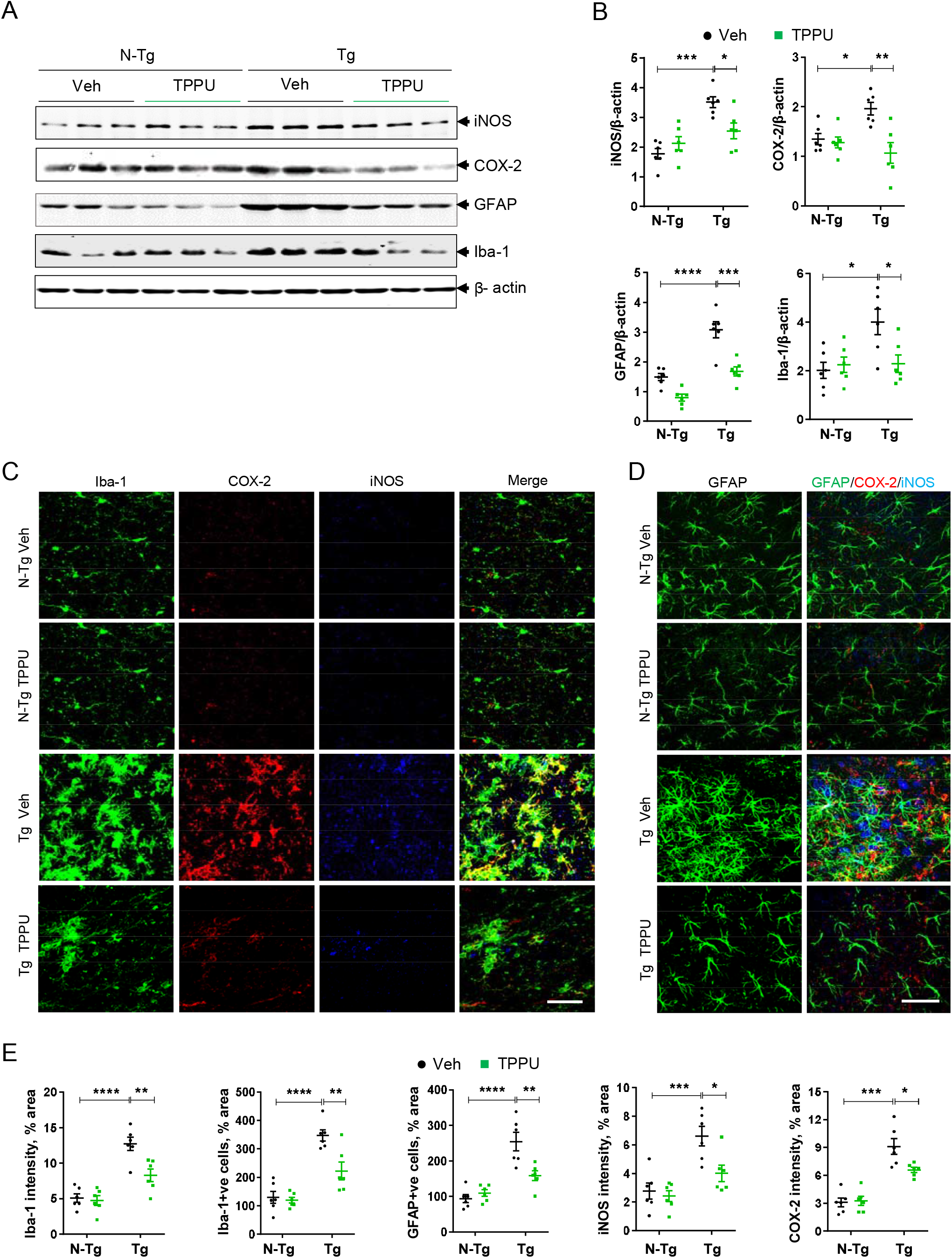
TPPU reduces neuroinflammatory markers and gliosis in Tg mice. (A) Representative Western blot of iNOS, COX-2, GFAP and Iba-1 in hippocampal samples of N-Tg and Tg mice treated with vehicle (Veh) or TPPU starting at 2 months for 2.5 months. β-actin was used as a control. (B) Quantification of (A). (C) Triple immunofluorescence staining of Iba-1 (green), COX-2 (red) and iNOS (blue) in hippocampus of above mice. Scale bar, 50 μm. (D) Immunofluorescence staining of GFAP (left) and merged panel of GFAP (green), COX-2 (red) and iNOS (blue) (right). Scale bar, 50 μm. (E) Quantification of Iba-1, iNOS and COX-2 intensities and Iba-1+ve and GFAP+ve cells in the hippocampus. Data are means ± SEM of six to eight mice per group. ****P < 0.0001, ***P < 0.001, **P < 0.01, *P < 0.05. Data were analyzed by two-way ANOVA with Bonferroni’s multiple comparison test.

### TPPU ameliorates Aβ pathology and functional impairment

Having demonstrated a significant role of TPPU in reversing AD-associated immune system dysfunction, we then asked whether sEH inhibition by TPPU may influence Aβ pathology. We stained the brain sections of 4.5 months and 6.5 months Tg mice treated with Veh or TPPU and quantified the Aβ plaque pathologies in the hippocampus (Fig. 5) and cortex (fig. S10). We observed modest 6E10-positive Aβ plaque deposition in 4.5 months Tg mice (Fig. 5A, Veh), which became more severe at 6.5 months (Fig. 5D, Veh). TPPU treatment led to significant reductions in the number, size, and intensities of Aβ plaques in both the 4.5 months (Fig. 5C) and the 6.5 months (Fig. 5F) groups. This was associated with reduced microglia activation surrounding Aβ plaques, marked by CD68 staining (Fig. 5, B-C, E-F). The same results were obtained when cortical samples were analyzed (fig. S10).

**Fig. 5.**
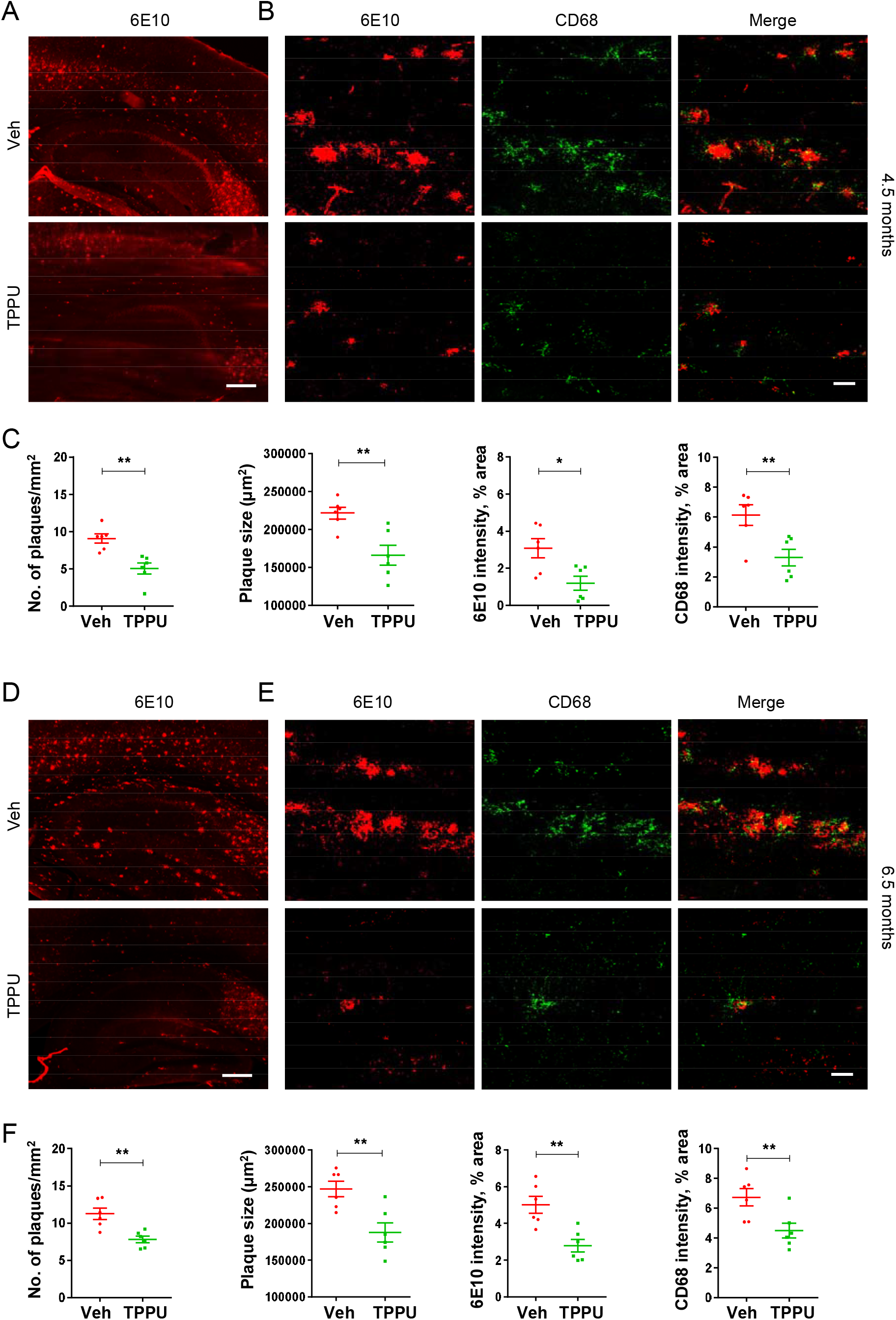
TPPU reduces A β burden in Tg mice. (A) Immunohistochemistry of 6E10 (lower magnification) in Tg mice treated with vehicle (Veh) or TPPU starting at 2 months for 2.5 months. Scale bar, 400 μm. (B) Higher magnification of hippocampus of Tg Veh or TPPU mice with double immunofluorescence staining of 6E10 (red) and CD68 (green). Scale bar, 200 μm. (C) Quantification of 6E10+ve plaque number, size and 6E10 and CD68 intensities in the hippocampus. (D-F) The same analysis and data presentation as (A-C) except mice treated with Veh or TPPU starting at 2 months for a duration of 4.5 months were analyzed. Data are means ± SEM of six to eight mice per group. **P < 0.01, *P < 0.05. Data were analyzed by Student’s *t*-test.

We next assessed the role of TPPU in rescue of neuronal phenotypes. Immunostaining with presynaptic protein synaptophysin (Syp) and high-resolution imaging of 6.5-month-old Veh or TPPU treated Tg mice and N-Tg controls observed significant reduction of Syn levels in area CA3 of hippocampus of Tg mice compared to N-Tg controls (Fig. 6A and quantified in B, Tg vs N-Tg, Veh). 4.5 months of TPPU treatment partially but significantly elevated the Syp expression (Fig. 6, A-B). LTP recordings of the Schaffer collateral pathway of the hippocampus revealed significant reductions in the Tg Veh group compared to N-Tg mice, and this phenotype was significantly improved in Tg TPPU mice (Fig. 6, C-D). Lastly, we evaluated the effect of TPPU in cognition using the novel object recognition (NOR) and fear conditioning (FC) paradigms (Fig. 6, E-F). The NOR assesses the hippocampus dependent long-term recognition memory by calculating the percent time spent with a novel object (object discrimination index, ODI). The Tg Veh mice displayed a significantly decreased ODI average, which was elevated significantly upon TPPU treatment (Fig. 6E). We further performed the FC paradigm to test hippocampal dependent (contextual test), and independent (cued test) associative learning (Fig. 6F). The four groups tested exhibited no differences in freezing percentage during the conditioning phase. During the context test, Tg vehicle mice displayed significantly decreased freezing percentage compared to N-Tg groups, suggesting an impaired contextual memory. Comparatively, the TPPU treated Tg mice exhibited significant increase in freezing frequency compared to the Tg Veh group. In addition, the percentage of freezing displayed post-cue was presented in the cued test was at similar levels between the groups. Thus, the Tg mice exhibit specific impairment in the hippocampus dependent contextual fear conditioning, and this phenotype is significantly improved by TPPU treatment. Taken together these results demonstrate that TPPU rescues synaptic deficits, LTP, and cognitive behaviors in Tg mice.

**Fig. 6.**
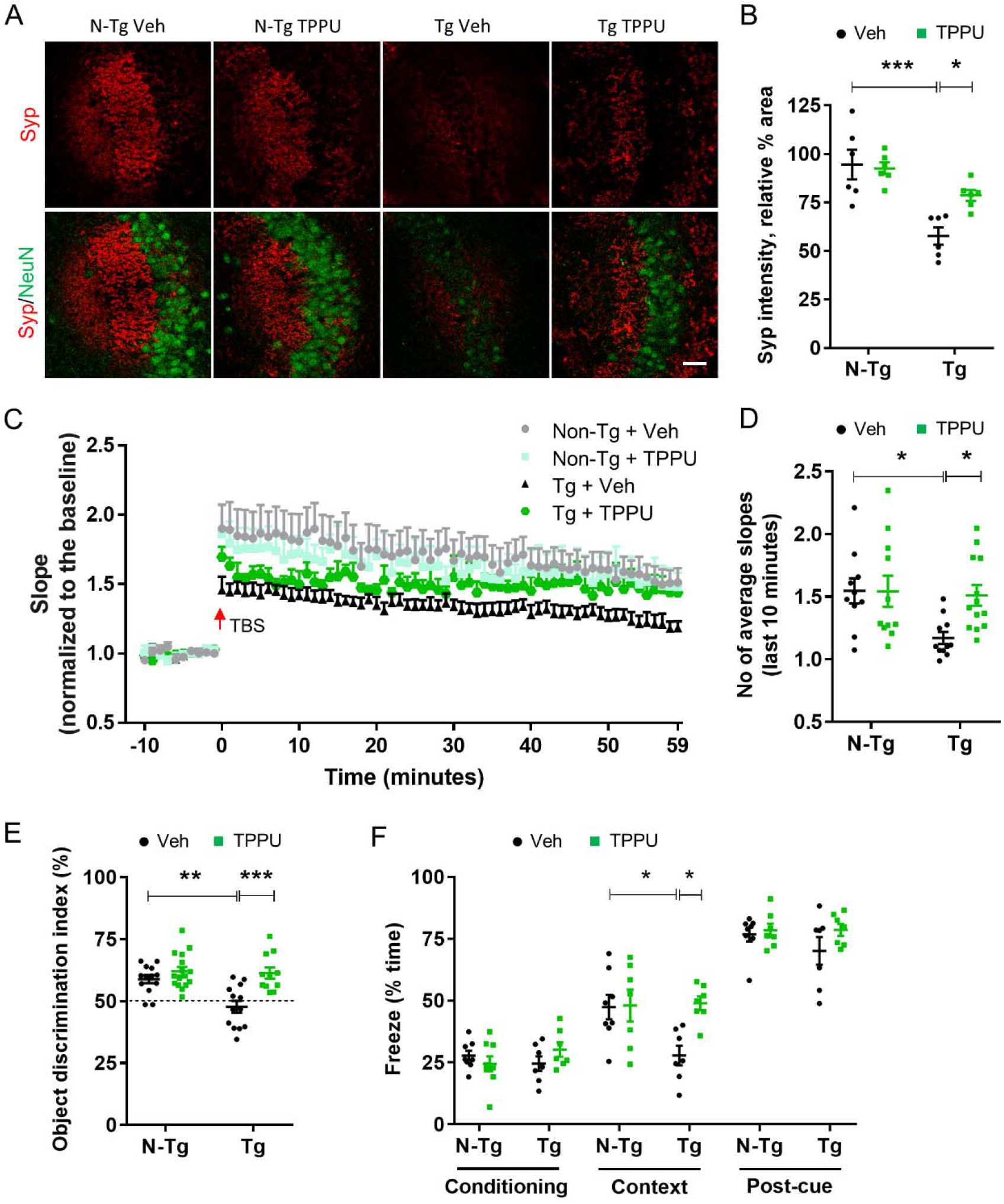
TPPU ameliorates synaptic deficits and cognitive function in Tg mice. (A) Representative images of synaptophysin (Syp) and NeuN co-immunostaining of area CA3 of hippocampus of N-Tg and Tg mice treated with vehicle (Veh) or TPPU starting at 2 months for 4.5 months. Scale bar, 100 μm. (B) Quantification of Syp intensity. (C) Slope of field excitatory postsynaptic potential (fEPSP) in response to theta burst stimulation delivered to the Schaffer collateral pathway from the above mice. Calibration: 2 mV, 5 ms. n = 10-13 sections from 5-6 animals per group/genotype. (D) No of average fEPSP slopes (last 10 minutes). (E) Object Discrimination Index of Veh- or TPPU-treated N-Tg or Tg mice in a novel object recognition test. The dotted line represents the 50% chance of random object exploration. (F) Percent of freezing in Veh- or TPPU-treated N-Tg or Tg mice in the Conditioning (left), Contextual (center) and cued (right) fear conditioning tests. Mice with 2.5 months Veh or TPPU treatment were subjected to behavior analysis. Values are expressed as mean ± SEM of either six to nine mice per group (A-D and F) or twelve to sixteen mice per group (E). Data were analyzed by two-way ANOVA with Bonferroni’s multiple comparison test. ***P < 0.001, **P < 0.01, *P < 0.05.

## Discussion

Using postmortem human brain samples, primary cell cultures and AD mouse models, we investigated the role of sEH in neuroinflammation and AD pathogenesis and tested the therapeutic effect of an orally bioavailable small molecule sEH inhibitor, TPPU. We found that the sEH levels are elevated in human AD brains and Aβ mouse models, the latter is well-correlated with significantly lower levels of EETs and EDPs. Pre-treatment with EET and TPPU prevent acute LPS-induced neuroinflammation. Long-term TPPU treatment at the onset of AD neuropathology is able to reverse microglia and astrocyte activation and immune pathway dysregulation at the molecular, cellular and functional levels, and these are associated with attenuated Aβ pathology and improved synaptic and cognitive function. Moreover, TPPU reinstates and positively correlates the EpFA levels in the Tg brain, supporting its brain penetration and target engagement.

Upregulation of sEH expression has been reported in CNS disorders such as depression (*26*), schizophrenia (*34*), and Lewy body dementia and Parkinson’s disease (*25*), and its inhibition has been shown to be beneficial in model systems. Lee et al (*27*) recently reported that sEH is upregulated in APP/PS1 mice. Our results clearly demonstrate that sEH is not only elevated in multiple transgenic AD mouse models, but also in human AD brains support the idea that it is a conserved and common feature in diseases with neuroinflammatory underpinning. Analysis of other molecules involved in ARA metabolism indicates partial activation of the ARA cascade. Interestingly, although the expression of *COX-2* and downstream *CYP4F8* are both elevated, only *EPHX2* but not CYP genes are altered. How this partial activation is achieved and whether this is also the case in other diseases is not understood. However, the data suggest that reduced EETs (and EDPs) are the result of their increased metabolism by sEH rather than insufficient conversion from ARA or release from phospholipids.

Our cell-type specific analysis demonstrates that astrocytes are the predominant cells expressing sEH where it is deregulated in AD conditions. This leads to diminished levels of EETs and EDPs and their anti-inflammatory activities on both astrocytes and microglia. Although no defined receptors have been identified for EETs, TRPV4 and G-proteins have been implicated in neuroinflammatory pathways downstream of EETs (*35*, *36*). Besides the astrocytes, sEH is known to be highly expressed in the vasculature where it mediates vascular inflammation and barrier function through both EETs and EDPs (*21*, *37*). Our expression analysis of sorted vascular endothelial cells revealed no appreciable differences in *Ephx2* expression between the Tg mice and N-Tg controls, arguing against a major contribution of vascular sEH in disease pathogenesis. Nevertheless, it remains possible that the overall therapeutic effect of TPPU is due to its inhibition of sEH in both astrocytes and vascular endothelia, and possibly other cell types. Interestingly, a recent report (*38*) showed that the liver sEH modulates depressive behaviors in mice through central-peripheral interactions. As such, although our finding that TPPU enters the brain where its levels correlate with EETs and EDPs supports a CNS intrinsic mechanism, we cannot exclude the possibility that TPPU could also exert its effect through inhibition of liver sEH. The availability of the *Ephx2* conditional allele (*38*) allows deciphering the cell-type specific effect and central-peripheral interactions.

EETs have been reported to act on multiple immune modulators, including the p38 MAP kinase, NF-κB, and STAT3 (*39*). EpFA are also known to stabilize mitochondria, reduce reactive oxygen species and shift the endoplasmic reticulum stress response from initiation of inflammation and cell death back towards maintaining cellular homeostasis (*40*). Thus, TPPU through EET and other EpFA could be acting on this pathway to reduce neuroinflammation. Of particular interest, we found in brain tissue as has been observed earlier in peripheral tissue that TPPU potently inhibits COX-2 expression (*26*), which has been widely implicated in AD. While the precise mechanism for this regulation remains to be established, it raises the intriguing possibility that TPPU’s anti-inflammatory activity may be conferred through inhibition of both sEH and COX-2.

Consistent with the above studies, our Nanostring analysis followed by qPCR validation revealed multiple immune and inflammatory response pathways are upregulated in the Tg mice and downregulated by TPPU. In addition, recent reports identified a signaling pathway whereby microglia mediated neuroinflammation, in a C1q-, IL-1α- and TNF-dependent manner, induces A1 astrocyte genes that are toxic to the neurons (*41*–*43*). Our gene expression analysis revealed that a number of A1 astrocyte genes (*Serping1*, *Gbp2*, *Srgn*, *H2T23*, *Psmb8*) were normalized by TPPU, suggesting that TPPU may reduce neuroinflammation via mitigating astrocytic activation, thus promoting neuronal survival. Additionally, the NLRP3 inflammasome activation has been implicated in producing harmful chronic inflammatory reactions and impairing microglial Aβ clearance and cognitive function in AD (*8*, *44*, *45*). Indeed, we found that *Nlrp3* and *Casp1* were both upregulated in Tg mice and downregulated by TPPU treatment. TPPU could suppress chronic neuroinflammation through NLRP3 dependent mechanisms.

We demonstrate that long-term TPPU treatment not only dampens glia reactivity but also ameliorates Aβ pathology and improves functional outcomes in Tg mice. Ample evidence documents that prolonged microglia activation leads to impaired Aβ phagocytosis and triggers the production of proinflammatory mediators, and its inhibition reverses these anomalies (*46*, *47*). Therefore, the observed reduction of Aβ pathology and improvement of neuronal function by TPPU could be the consequences of glia normalization. Additionally, microglia, through complement-dependent mechanisms, has been shown to mediate synapse elimination (*48*, *49*). Since we observed robust rescue of aberrant *C1q* and *C3* expression in Tg mice by TPPU (Fig. 3), the augmented synaptic protein expression and behavioral performance could be attributed by improved synapse maintenance.

Several classes of sEH inhibitors have been developed (*50*). Overall, they are well-tolerated in preclinical studies, establishing the large safety window for sEH targeting. Among these, TPPU is widely used as a tool compound because of its superior potency, specificity, and pharmacokinetics (*23*, *51*–*54*). Of particular interest, a Phase 1 trial was recently initiated to test a TPPU analog (EC5026) for neuropathic pain (https://www.prnewswire.com/news-releases/eicosis-announces-first-subject-dosed-in-phase-1a-clinical-trial-of-ec5026-300971946.html). We report here that long-term administration of TPPU results in significant brain retention where it engages its target and affords beneficial effect in a mouse model of AD. These features make TPPU an attractive lead candidate for the treatment of AD and possibly other neurodegenerative diseases.

## Materials and Methods

### Study design

The goal of this study was to establish evidence, using multiple model systems, that inhibition of the soluble epoxide hydrolase via TPPU is a viable approach to modify disease progression in AD. In the setting of experiment, one individual would randomize the animals, plates, and slides, and another would analyze them. The minimum sample size for all experiments was held at six mice per group based on the design of previous studies (*11*). To improve our power, and thus our ability to statistically detect smaller effects, many of our analyses included more mice per group. Further experimental details and protocols of each model, including animal care/handling and the number of biological/technical replicates, are in this section, and in Figure Legends.

### Human subjects

Postmortem brain tissues were provided by the University of Pennsylvania Center for Neurodegenerative Disease Research (CNDR). Informed consent was obtained from all subjects. The demographic data can be found in Supplementary Table 2. Influence of sex, gender identity or both on the study results was not the objective of the study. It was not analyzed due to small sample size.

### Mice and treatment

The C57BL/6, and 5xFAD mice were obtained from the Jackson Laboratory (Bar Harbor, ME). APP^NLGF^ mice were obtained from RIKEN (*28*). Mice were housed 4-5 per cage in a pathogen free mouse facility with ad libitum access to food and water on a 12 hr light/dark cycle. Male and female mice at approximately equal ratio were used unless otherwise specified. All procedures were performed in accordance with NIH guidelines and approval of the Baylor College of Medicine Institutional Animal Care and Use Committee (IACUC).

In the acute TPPU regimen, ten- to twelve-week-old C57BL/6 mice received 1^st^ dose of TPPU (3 mg/kg) via oral gavage 24 h before co-treatment of LPS (3 mg/kg, i.p.) and TPPU (2^nd^ dose). Eighteen-hour post co-treatment, mice were sacrificed for analysis. The sEH inhibitor TPPU was synthesized as previously described (*23*). TPPU was dissolved either in DMSO for in vitro and ex vivo treatment or in 10% polyethylene glycol 400 (PEG400, Fisher) for in vivo treatment. The vehicle mice received oral gavage treatment of 1% PEG400. Mice were perfused with saline before collecting brain for biochemical analysis.

### Primary microglia and astrocyte cultures

Primary glia cultures were prepared as described previously (*48*). In brief, mouse cortices and hippocampi were isolated from newborn pups (P0-P1) in dissection medium (HBSS with 10 mM HEPES, 1% v/v Pen/Strep) and cut into small pieces. Tissue was digested with 2.5% trypsin at 37°C for 15 min before trypsin inhibitor (1 mg/ml) was added. Next, tissue was centrifuged for 5 min at 1500 rpm, triturated, and resuspended in DMEM medium with 10% FBS. Cells were plated onto poly-D-lysine (PDL)-coated T-75 flasks at 50,000 cells/cm^2^ to generate mixed glial cultures. When confluent, microglia were separated by tapping the flasks against table and collecting the floating cells in media. Microglia cells were then seeded at 50,000 cells/cm^2^ and cultured for another day in PDL-coated 12-well plates for mRNA assays or on coverslips for staining. After collecting microglia cells, remaining cells (mostly astrocytes) were trypsinized and seeded at 40,000 cells/cm^2^ and cultured for another two days in PDL-coated plated for mRNA assays or immunocytochemistry.

### RNA extraction and expression analysis

Total RNA was extracted from cells or human or mouse brain tissues using RNeasy Mini kit (Qiagen, 74106). Reverse transcription was carried out using iScript Reverse Transcription Supermix (Bio-Rad, 1708840). The qPCR analyses were performed using SYBR Green PCR master mix (Bio-Rad) on a CFX384 Touch Real-Time PCR Detection System. Primer sequences can be found in Supplementary Table 3.

For Nanostring analysis, RNA was isolated from 4.5-month-old mouse hippocampus and 770 transcripts were quantified with the Nanostring nCounter multiplexed target platform using the Mouse Neuroinflammation panel (https://www.nanostring.com). nCounts of mRNA transcripts were normalized using the geometric means of 10 housekeeping genes (*Csnk2a2*, *Ccdc127*, *Xpnpep1*, *Lars*, *Supt7l*, *Tada2b*, *Aars*, *Mto1*, *Tbp*, and *Fam104a*) and analyzed using nSolver 4.0 and the Advanced Analysis 2.0 plugin. Fold-change expression and p-values were calculated by linear regression analysis using negative binomial or log-linear models. P-values were corrected for multiple comparisons using the Benjamini-Yekutieli method. Volcano plots of differential expression data were plotted using the −log10 (p-value) and log2 fold-change using the Graphpad prism. Gene ontology enrichment analysis was performed using https://www.innatedb.com/. Heatmaps were constructed using Graphpad prism.

### Griess assay

Griess assay was performed, as described previously (*55*). Briefly, standards in triplicate were used for each plate. Standard mix was made up of 1 ml media and 1 μl sodium nitrite and added to wells in volumes increasing by 5 μl from 0-35 μl. Media was added to each well to bring volume up to a total of 100 μl. Sample supernatant was added in duplicate to remaining wells in 100 μl. Then, 100 μl of Griess reagent (Sigma) was added to each well and incubated at RT for 20 minutes. Absorbance at 540 nm was detected in a Synergy 2 Multi-Detection Microplate Reader.

### P-glycoprotein substrate evaluation

Caco-2 cells were diluted to 6.86×10^5^ cells/mL with culture medium and 50 μl of cell suspension were dispensed into the filter well of the 96-well HTS Transwell plate. Cells were cultivated for 14-18 days in a cell culture incubator at 37°C, 5% CO_2_, 95% relative humidity. Electrical resistance was measured across the monolayer by using Millicell Epithelial Volt-Ohm measuring system. "TEER of each well is calculated by the equation-TEER value (ohm•cm^2^) = TEER measurement (ohms) x Area of membrane (cm^2^). The TEER value of each well should be greater than 230 ohms•cm^2^. Digoxin was used as the reference substrate of P-gp. Propranolol was used as the high permeability marker. To determine the rate of drug transport in the apical to basolateral direction, working solutions containing TPPU was added to the Transwell insert (apical compartment). To determine the rate of drug transport in the basolateral to apical direction, working solutions containing TPPU was added to each well of the receiver plate. To determine the rate of drug transport in the presence of the P-gp inhibitor, verapamil the known inhibitor of Pgp, was added to both apical and basolateral compartments at a final concentration of 100 μM, followed by incubation at 37 °C for 2 hours. Next, samples from apical and basolateral wells were transferred to a new 96-well plate and cold acetonitrile containing appropriate internal standards (IS) were added into each well of the plate(s). Samples were analyzed by an LC-MS/MS. Percent parent compounds remaining at each time point are estimated by determining the peak area ratios from extracted ion chromatograms. The apparent permeability coefficient (P_app_), in units of centimeter per second, using the following equation: P_app_ = (VA×[drug]acceptor)/(Area×Time×[drug]initial, donor), where VA is the volume (in ml) in the acceptor well, area is the surface area of the membrane (0.143 cm^2^ for Transwell-96 Well Permeable Supports), and time is the total transport time in seconds. The efflux ratio was determined using the following equation: Efflux Ratio=P_app_(B-A)/P_app_(A-B), where P_app_ (B-A) indicates the apparent permeability coefficient in basolateral to apical direction, and P_app_ (A-B) indicates the apparent permeability coefficient in apical to basolateral direction. The recovery can be determined using the following equation: Recovery%=(VA×[drug]acceptor+VD×[drug]donor)/(VD×[drug]initial, donor), where VA is the volume (in ml) in the acceptor well (0.235 ml for Ap→Bl flux, and 0.075 ml for Bl→Ap), VD is the volume (in ml) in the donor well (0.075 ml for Ap→Bl flux, and 0.235 ml for Bl→Ap).

### Cell-type purification and FACS sorting

Mice were perfused with ice-cold PBS, adult mouse brains (whole brain minus cerebellum) were chopped and resuspended in 2.5 mls of HBSS w/o Ca^2+^, and w/o Mg^2+^ containing activated papain and DNase. Cell type purification and FACS sorting were done as described (*29*). Briefly, brains were incubated at 37°C, then triturated 4 times with a fire-polished glass Pasteur pipet. Next, samples were mixed with HBSS+ (HBSS + 0.5% BSA, 2mM EDTA) and centrifuged for 5 min. The pellet was resuspended in 1000 ml of HBSS+ and centrifuged for 15 sec at room temperature. The supernatant was collected and was filtered through cell strainer (BD SKU 352340) and centrifuged for 5 min at 300 g at 4°C. To remove myelin, the Miltenyi myelin removal beads were used according to the manufacturer’s instructions (Miltenyi, 130-096-733). After that, cells were centrifuged at 300xG for 5 min at 4°C. Next, the cells were resuspended in 1ml HBSS+ solution and passed through a LS column. The total effluent was then centrifuged for 5 min at 300 g at 4°C to pellet the cells. For the antibody staining, the cells were resuspended in HBSS+ solution and then stained for with CD45-BV421 (BD, 563890), CD11b-FITC (BD, 553310) for microglia, ACSA-2-APC (Miltenyi, 130-102-315) for astrocytes, and cell viability blue fluorescent dye (Invitrogen, L23105). After FACS sorting, the cells were collected in Eppendorf tubes, centrifuged at 1500 rpm for 5 min, and resuspended in RLT buffer containing 1% BME for future qPCR analysis. The mRNA was extracted using the QIAGEN RNAEasy Micro kit (QIAGEN, 74004).

### Immunostaining and quantification

Cells were fixed with 4% paraformaldehyde in 1X PBS for 15 min and processed for immunocytochemistry, as described previously (*56*). First, nonspecific sites were blocked with 0.2% bovine serum albumin, 0.5% Triton X-100, and 0.05% Tween 20 in PBS for 1 h at room temperature. Cells were then incubated with different primary antibodies: sEH (1:1000, Dr. Bruce Hammock lab (*26*)); GFAP (1:1000, Millipore); iNOS (1:500, Thermo Fisher); Iba-1 (1:1000, Wako) and COX-2 (1:1000, Thermo Fisher) at 4°C overnight. Appropriate secondary antibodies (Alexa Fluor 488 or 555 or 647, Invitrogen) were used followed by incubation with DAPI to stain the nucleus. The coverslip-containing stained cells were washed twice with PBS and mounted on slides. After immunofluorescent staining, five individual areas from each coverslip were imaged using 20x magnification under a Leica TCS confocal microscope. Mean intensity of fluorescence and number of immunoreactive cells were quantified using the Image J software (NIH).

Immunohistochemistry was performed on free-floating microtome-cut sections (30 μm in thickness), as described previously (*55*). Briefly, mice brains were post fixed in 4% paraformaldehyde overnight at 4°C and transferred to 30% sucrose solution following perfusion with saline. Sections were incubated with different antibodies: sEH (1:1000, Dr. Bruce Hammock lab (*26*)); GFAP (1:1000, Millipore); iNOS (1:250, Thermo Fisher); Iba-1 (1:800, Wako or 1:500, Novus Biologicals), COX-2 (1:500, Thermo Fisher), CD68 (1:500, BioLegend), 6E10 (1:1000, BioLegend). Appropriate secondary antibodies (Alexa Fluor 488 or 594 or 647, Invitrogen) were used followed by incubation with DAPI. A total of three to four sections per brain containing the hippocampus and cortex and five to seven mice per group were stained with antibodies as mentioned above. After immunofluorescence staining, confocal images were captured and mean intensity of fluorescence and number of immunoreactive cells were quantified using the Image J software (NIH). For quantification of 6E10 in the mouse cortex and hippocampus, sections were scanned using an EVOS FL Auto system. Images were then analyzed by ImageJ and background was subtracted by the software for fluorescence images before quantification.

### Western blotting

Cells or brain tissues were collected and resuspended in modified radioimmunoprecipitation (RIPA) assay buffer containing protease and phosphatase inhibitor mixture. Cell suspensions were sonicated after resuspension, whereas mouse brain tissues were homogenized, sonicated, and then centrifuged at 14,000 × g for 45 min at 4°C, as described previously (*57*). Briefly, protein concentrations were estimated using a BCA kit (Thermo Fisher). Lysates were separated on 7.5%–15% SDS-polyacrylamide electrophoresis gels (Bio-Rad). After the separation, proteins were transferred to a nitrocellulose membrane, and nonspecific binding sites were blocked by treating with either Odyssey blocking buffer (LI-COR) or TBS with 5% bovine serum albumin (BSA) followed by antibody incubation: sEH (1:300, Santa Cruz or 1:1000, Dr. Bruce Hammock’s lab (*26*)); iNOS (1:1000, Cell signaling); GFAP (1:15000, EMD Millipore); Iba-1 (1:1200, Wako); Cox-2 (1:500, Thermo Fisher); and β-actin (1:10,000, Sigma). Secondary IR-680-conjugated goat anti-mouse or goat anti-rabbit (1:10,000; Molecular Probes), IRDye 800-conjugated donkey anti-rabbit or donkey anti goat (1:10,000, Rockland, PA, USA) were used. Western blot images were captured with a LI-COR Odyssey machine (LI-COR). The western blot bands were quantified using ImageJ software (NIH).

### Behavioral analysis

The novel object recognition (NOR) protocol included three phases; habituation phase, a training phase and an object recognition phase. The habitation phase included of one session, 5 minutes in length, in which the animals were allowed to freely explore a small Plexiglas arena (measuring 22cm × 44cm) that was utilized in the training and testing phase. One day after habituation the animals underwent training. During the training phase, the animals were placed in the same arena with the addition of two identical objects. The animals were allowed to freely explore the objects for 5 minutes. 24 hours after the training phase, the test phase was initiated. During the testing phase, the animal was placed in the same arena with one object previously explored in the training phase, the familiar object, and one novel object differing in color and shape, but sharing a common size and volume. The animals were allowed to freely explore the objects for 5 minutes. ANY-Maze software was used to measure time spent exploring each object. Exploration of an object was defined by head orientation directed toward the object or physical contact with the object. The object discrimination ratio (ODR) was calculated by the following formula: ODR= (Time exploring specified object)/ (time exploring novel object+ time exploring familiar object) × 100. The fear conditioning protocol involved a training phase, context test, and a cued test as previously described (*58*). During the training phase the mice were placed in the training chamber and allowed to freely explore the environment. At 3 minutes, an 80-dB white noise was presented (auditory conditioned stimulus (CS)) for 30 seconds. During the last 2 seconds of the auditory stimulus, the unconditioned stimulus (US), a foot shock (0.8 mA, 2 seconds), was administered. The CS and US were then presented a second time at the 5-minute mark of the training procedure. After the second presentation of the US, the mice stayed in the training chamber for an additional 2 minutes without additional stimulations. The animals were returned to their original housing cages. 24 hours after the training procedure, the context test was performed. The mice were returned to the same training chamber consisting of the same context as the first procedure (same geometric shape of chamber, lights, scents and auditory sounds) for 3 minutes with no presentations of US or CS. One hour later, the cue test is performed. The cue test chamber consisted of a different geometric shape, flooring, light brightness and scent compared to the previous chamber used for training. After 3 minutes in the chamber, the auditory stimulus was presented for 3 minutes. The software, FreezeFrame3 and FreezeView (San Diego Instruments) was used to record and analyze the percent freezing in each trial.

### Electrophysiology

Field recordings of Schaffer collateral LTP was performed as described before (*59*). Briefly, brains were isolated from 6.5-7-month-old mice and cut into 400 mM slices on a vibratome. Hippocampal slices were incubated for 1 h at room temperature and then transferred to a heated recording chamber filled with recording ACSF (125 mM NaCl, 2.5 mM KCl, 1.25 mM NaH_2_PO_4_, 25 mM NaHCO_3_, 1 mM MgCl_2_, 2 mM CaCl_2_, and 10 mM glucose, saturated with 95% O_2_ and 5% CO_2_) maintained at 32°C. Stimulation of Schaffer collaterals from the CA3 region was performed with bipolar electrodes, while borosilicate glass capillary pipettes filled with recording ACSF (resistances of 2–3.5 MΩ) were used to record field excitatory postsynaptic potentials (fEPSPs) in the CA1 region. Signals were amplified using a MultiClamp 700 B amplifier (Axon), digitized using a Digidata 1440A (Axon) with a 2 kHz low pass filter and a 3 Hz high pass filter and then captured and stored using Clampex 10.4 software (Axon) for offline data analysis. The genotypes and treatment groups were blinded to the experimenter. For each experiment 10-13 sections from 5-6 animals per group/genotype were used.

### Measurement of brain and plasma drug concentrations

5xFAD mice were treated with TPPU (3 mg/kg) in drinking water for 4 months and then euthanized. Plasma and whole-brain homogenates were extracted and subjected to LC-MS/MS analysis of TPPU and oxylipin on a 4000 Qtrap LC-MS/MS instrument (Applied Biosystems Instrument Corporation). For drug analysis in plasma, 10 μl of plasma samples were transferred to 1.5 μl Eppendorf tubes containing 90 μl EDTA solution (0.1% EDTA and 0.1% acetic acid), spiked with 10μl of 1μg/ml TPPU-d3 in methanol, and subsequently subjected to liquid-liquid extraction by ethyl acetate (200 μl) twice (*54*). TPPU-d3 was added in each sample as an internal standard solution. The collected extraction solutions were dried using a speed vacuum concentrator, reconstituted in 50 μl of 100 nM CUDA in methanol, and ready for LC-MS/MS analysis. TPPU and oxylipin in brain tissues were analyzed simultaneously by a modified LC-MS/MS method(*60*) to including MRM transition of TPPU. Tissues (~50 mg) were homogenized in ice cold methanol with 0.1% BHT and 0.1% acetic acid. The homogenates were spiked with 10μl of internal standard solution (mixture of deuterated compounds) and stored at −80°C for 20hr.

After that, the homogenates were extracted using solid-phase extraction (Oasis-HLB Cartridge, Waters). The extracted samples were then collected, dried and reconstituted in 50μl of 200 nM CUDA in methanol. The analytes were then detected by the modified LC-MS/MS method.

### Quantification and statistical analysis

All data were analyzed with GraphPad Prism v.7.04 and presented as mean ± SEM (*P < 0.05, **P < 0.01, ***P < 0.001 and ****P < 0.0001). For simple comparisons, Student’s t test was used. For multiple comparisons, ANOVA followed by the appropriate post hoc testing was utilized and is specified for each experiment in the figure legends. The statistical tests used for human data expression analysis is specified in the human data analysis methods section. All samples or animals were included in the statistical analysis unless otherwise specified.

## Acknowledgements

We are grateful to V. Lee and J. Trojanowski (University of Pennsylvania) for providing postmortem brain tissues and T. Saito and T. Saido (RIKEN Brain Science Institute) for the APP^NLGF^ knock-in mice. We thank Bianca Contreras and N. Aithmitti for expert technical assistance, B. Wang and E. Roy for help with organotypic culture and Nanostring experiments and members of the Zheng laboratory for insightful discussions. We acknowledge support from the Genomic and RNA Profiling Core and the Cytometry and Cell Sorting Core at Baylor College of Medicine for Nanostring and FACS analyses, respectively.

## Author Contributions

AG and HZ designed the overall study; AG performed all in vitro and in vivo assays and associated biochemical and immunohistochemical experiments and data analysis, with technical assistance from MC and NEP; MC and FC performed behavioral tests and LTP recording and associated data analysis, respectively; SHH and BDH provided TPPU and advised on the related experiments; DW performed mass spec analysis of oxylipins; AG and HZ wrote the manuscript and all authors read, provided input and approved the manuscript.

## Data and materials availability

All data used for this study are included in the main manuscript or supplemental materials.

