## Supplemental figures and tables for "Epoxy fatty acid dysregulation and neuroinflammation in Alzheimer’s disease is resolved by a soluble epoxide hydrolase inhibitor"

**Supplementary Materials**

Fig. S1. Expression of genes related to arachidonic acid (ARA) pathway in transgenic AD mouse models.

Fig. S2. LPS induces sEH expression in astrocytes.

Fig. S3. TPPU attenuates LPS-induced sEH expression and inflammation in primary astrocytes.

Fig. S4. TPPU fails to mitigate inflammation in primary microglia.

Fig. S5. EET attenuates LPS-induced inflammation in primary microglia.

Fig. S6. EET attenuates LPS-induced inflammation in organotypic hippocampal slice cultures.

Fig. S7. TPPU mitigates LPS-induced inflammation in the hippocampus of C57BL/6 mice.

Fig. S8. TPPU mitigates LPS-induced inflammation in C57BL/6 mice.

Fig. S9. TPPU reduces neuroinflammation in Tg mice.

Fig. S10. TPPU reduces Aβ burden in Tg mice.

Table S1. P-glycoprotein (P-gp) substrate evaluation of TPPU in Caco-2 cell line.

Table S2. Human demographics related to “Human subjects” in methods.

Table S3. Primer sequences for qPCR. Related to “RNA extraction and quantitative PCR (qPCR)”.

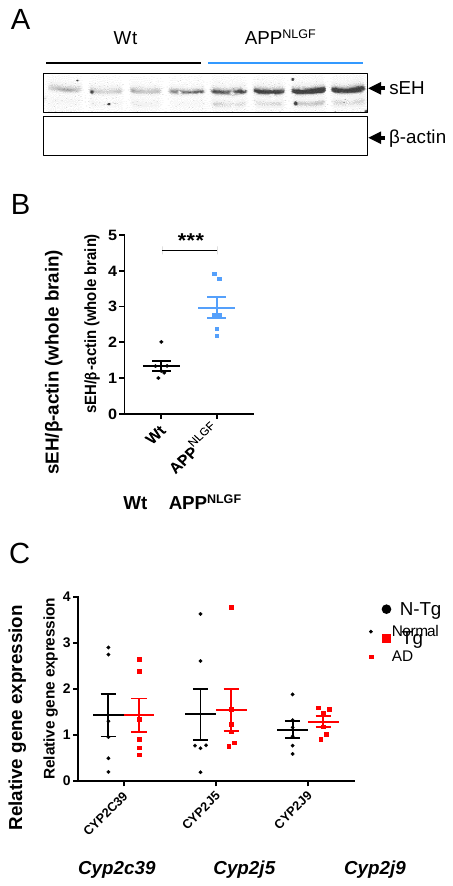

**Fig. S1.** **Expression of genes related to arachidonic acid (ARA) pathway in transgenic AD mouse models.** (A) Representative western blot illustrating the expression of sEH in APP^NLGF^ mice at 6.5 months of age. (B) Scattered dot plot showing mean Western blot sEH/β-actin ratio in APP^NLGF^ mice. (C) qPCR analysis of different subunits of cytochrome P450 (Cyp) in Tg mice at 4.5 months of age. Values are expressed as means ± SEM of six to eight mice per group. Data were analyzed by unpaired Student’s *t*-test. ***P < 0.001, **P < 0.01, *P < 0.05.

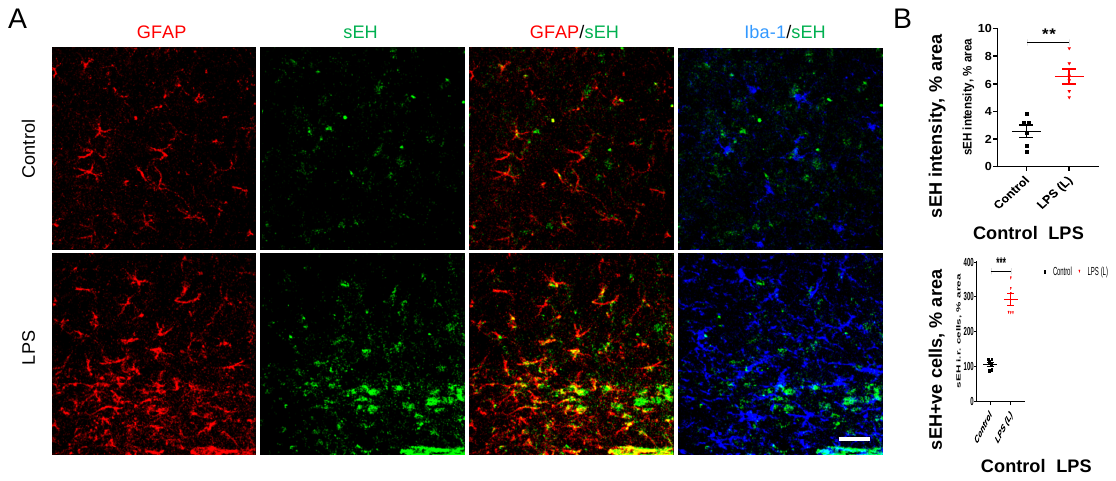
**Fig. S2.** **LPS induces sEH expression in astrocytes.** (A) Eighteen-hour post LPS treatment, C57BL/6 mice were euthanized, and hippocampal sections were processed for immunohistochemistry of GFAP (red) and sEH (green). Scale bar, 50 μm. (B) sEH intensity (top) and sEH +ve cells (bottom) in the hippocampus. Values are expressed as means ± SEM of six mice per group. Data were analyzed by unpaired Student’s *t*-test. ***P < 0.001, **P < 0.01.

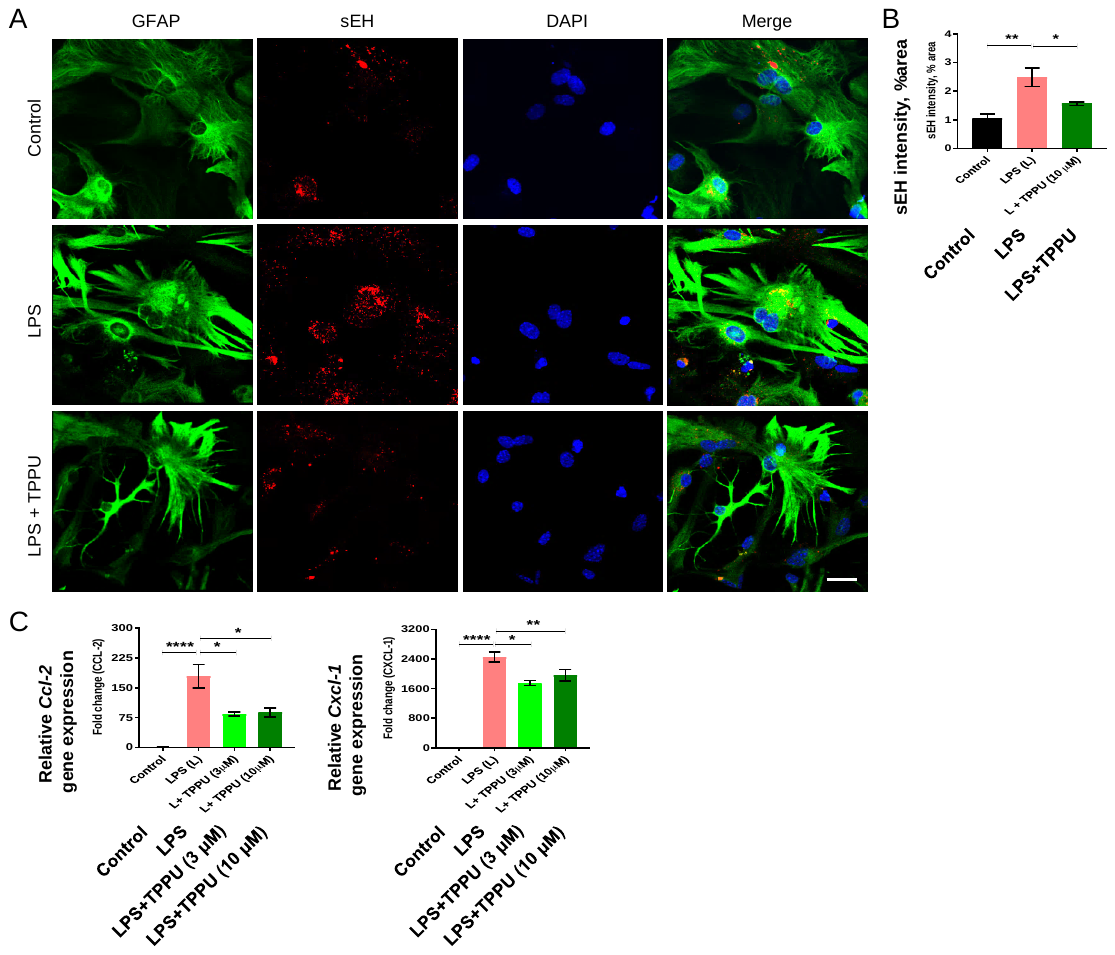

**Fig. S3.** **TPPU attenuates LPS-induced sEH expression and inflammation in primary astrocytes.** Primary astrocytes were pretreated with multiple doses of TPPU (3 μM and 10 μM) for 30 minutes followed by LPS treatment (100 ng/ml) for 24h. (A) Double labeled immunofluorescence of GFAP (green) and sEH (red) in primary astrocytes. DAPI stains (blue) nucleus in the merge panel. Scale bar, 100 μm. (B) Quantification of sEH intensity. (C) qPCR analysis of mRNA expression of *Ccl-2* and *Cxcl-1*. Data are means ± SEM of three independent experiments. ****P < 0.0001, ***P < 0.001, **P < 0.01, *P < 0.05. Data were analyzed by one-way ANOVA with Tukey’s multiple comparison test.

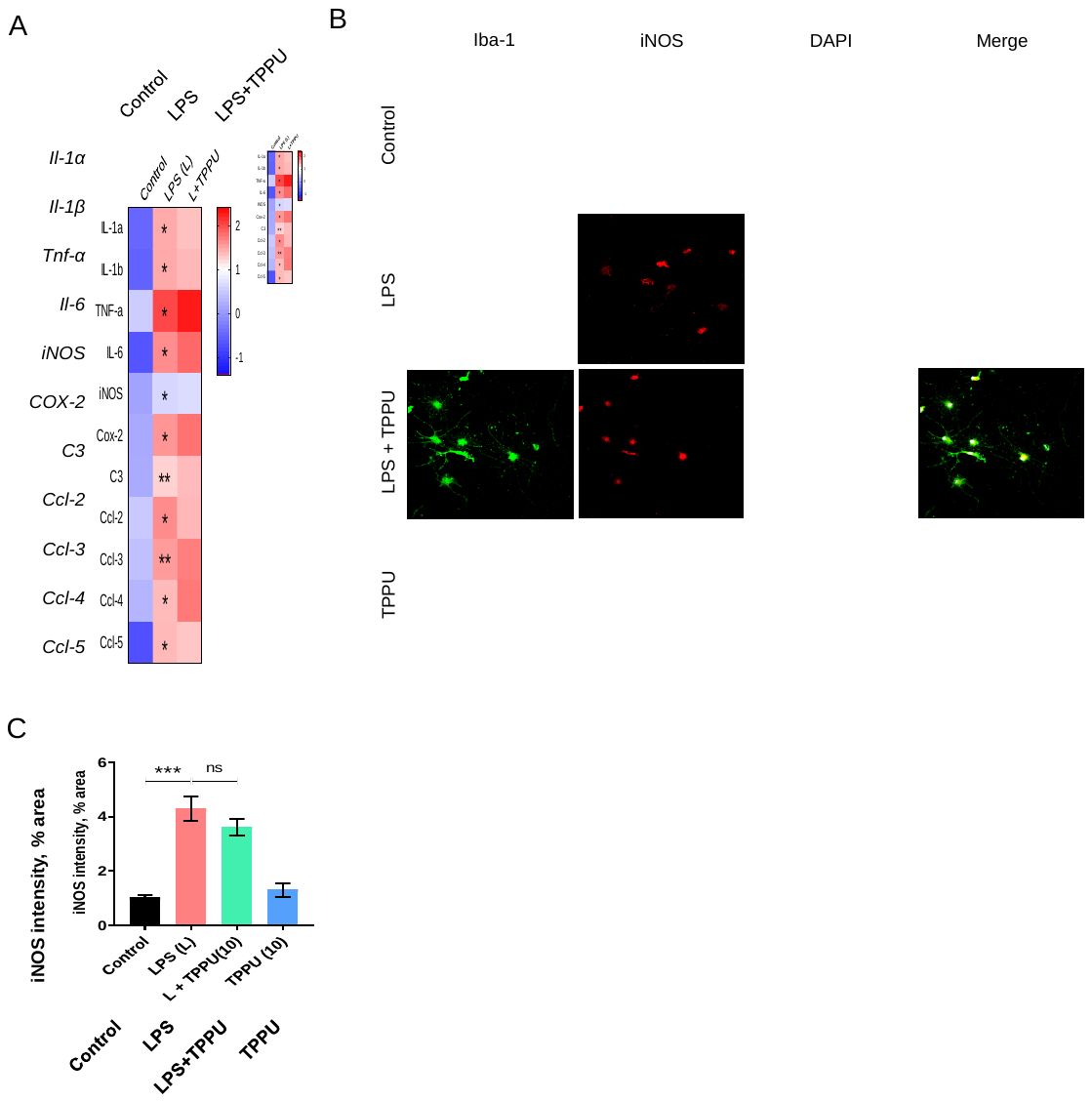

**Fig. S4.** **TPPU fails to mitigate inflammation in primary microglia.** Primary microglia were pretreated with 10 μM TPPU for 30 minutes followed by LPS treatment (100 ng/ml) for 24h. (A) Heat map showing qPCR analysis of mRNA expression of inflammatory genes. The asterisks in LPS column represent the comparison between control and LPS. (B) Immunocytochemistry of Iba-1 (green) and iNOS (red). DAPI stains the nucleus (white) in the merge panel. Scale bar, 75 μm. (C) iNOS intensity in primary microglia. Data are means ± SEM of three independent experiments. ***P < 0.001, **P < 0.01, *P < 0.05, ns= not significant. Data were analyzed by one-way ANOVA with Tukey’s multiple comparison test.

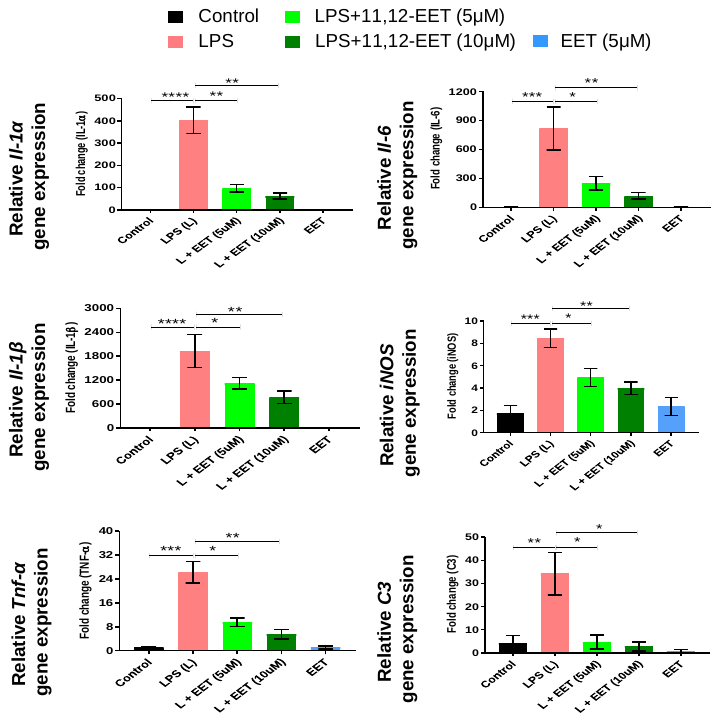

**Fig. S5.** **EET attenuates LPS-induced inflammation in primary microglia.** Primary microglia were pretreated with 5 μM or 10 μM of 11,12-EET for 30 minutes followed by LPS treatment (100 ng/ml) for 24 h. qPCR analysis of mRNA expression of *Il-1α*, *Il-1β*, *Tnf-α*, *Il-6*, *iNOS* and *C3*. Data are means ± SEM of three independent experiments. ****P < 0.0001, ***P < 0.001, **P < 0.01, *P < 0.05. Data were analyzed by one-way ANOVA with Tukey’s multiple comparison test.

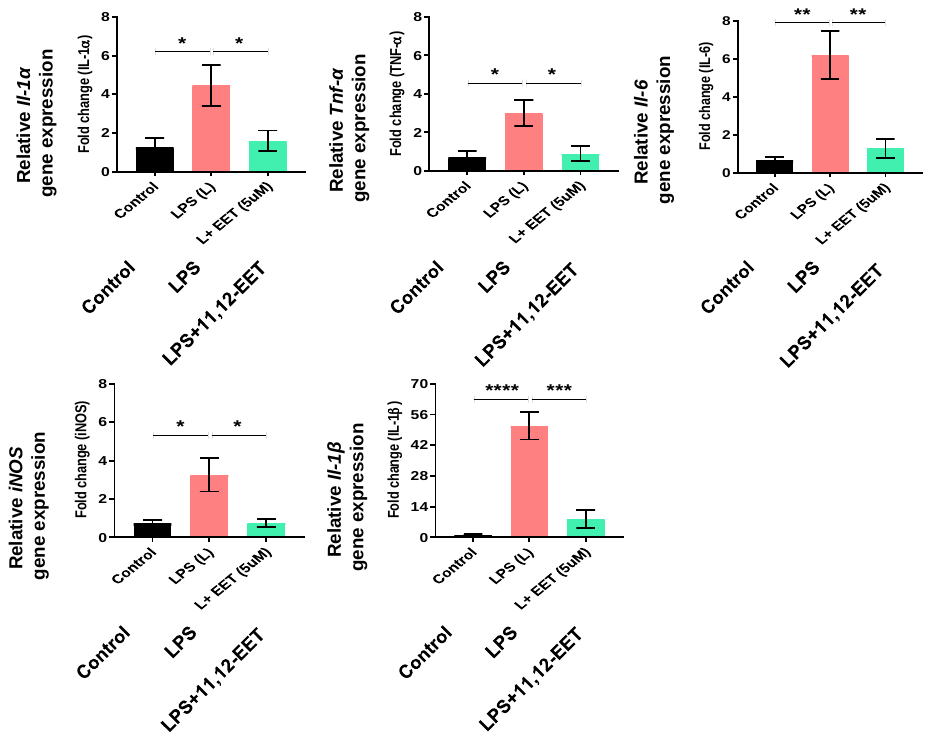

**Fig. S6.** **EET attenuates LPS-induced inflammation in organotypic hippocampal slice cultures.** Hippocampal slices from C57BL/6 mice were pretreated with 11,12-EET (5 μM) for 30 minutes followed by LPS treatment (100 ng/ml) for 24h. qPCR analysis of mRNA expression of pro-inflammatory molecules. Data are means ± SEM of three independent experiments. ****P < 0.0001, ***P < 0.001, **P < 0.01, *P < 0.05. Data were analyzed by one-way ANOVA with Tukey’s multiple comparison test.

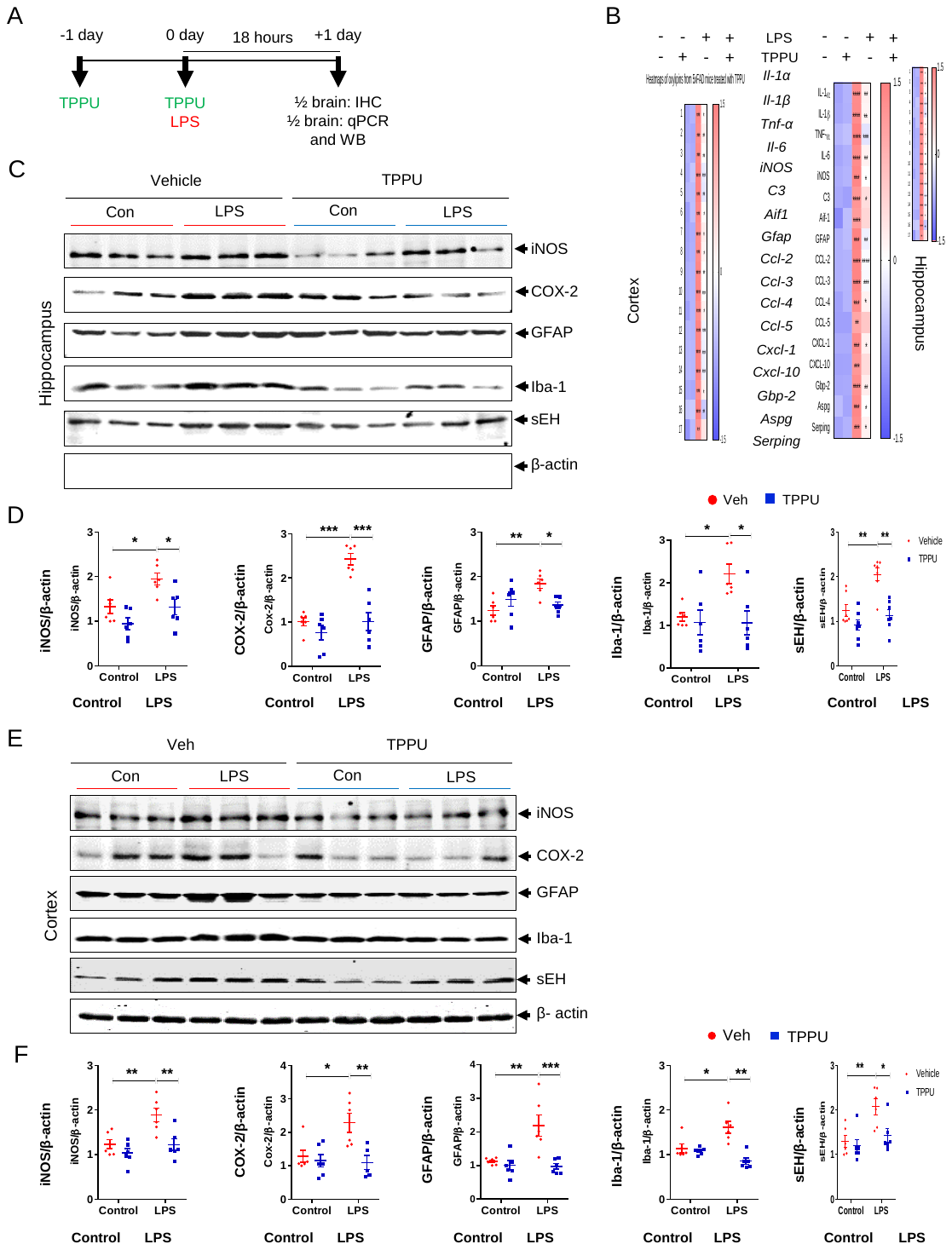

**Fig. S7.** **TPPU mitigates LPS-induced inflammation in the hippocampus of C57BL/6 mice.** (A) Schematic diagram showing TPPU treatment in C57BL/6 mice. Mice were pretreated with TPPU (3 mg/kg, oral-gavage) 1 day before (-1d) LPS administration. On day 0 (0d) mice were co-treated with LPS (3 mg/kg, i.p.) and TPPU (3 mg/kg, oral gavage). Eighteen-hour post co-treatment, mice were euthanized and processed for biochemical analysis. One hemisphere of brain was processed for IHC and the other hemisphere was used for qPCR and WB. (B) Heat map visualization of qPCR analysis of mRNA expression in Cortex (left) and in Hippocampus (right). The asterisks in LPS (+) TPPU (-) column represent significant changes vs LPS (-) TPPU (-). The asterisks in LPS (+) TPPU (+) represent significant change vs LPS (+) TPPU (-). (C) Representative Western blot illustrating the expression iNOS, COX-2, GFAP, Iba-1 and sEH in hippocampus. (D) Scattered dot plot showing mean Western blot protein expression over β-actin in hippocampus. (E) Representative Western blot illustrating the expression iNOS, COX-2, GFAP, Iba-1 and sEH in cortex. (F) Scattered dot plot showing mean Western blot protein expression over β-actin in cortex. Data are means ± SEM of six to eight mice per group. ***P < 0.001, **P < 0.01, *P < 0.05. Data were analyzed by one-way ANOVA with Tukey’s multiple comparison test.

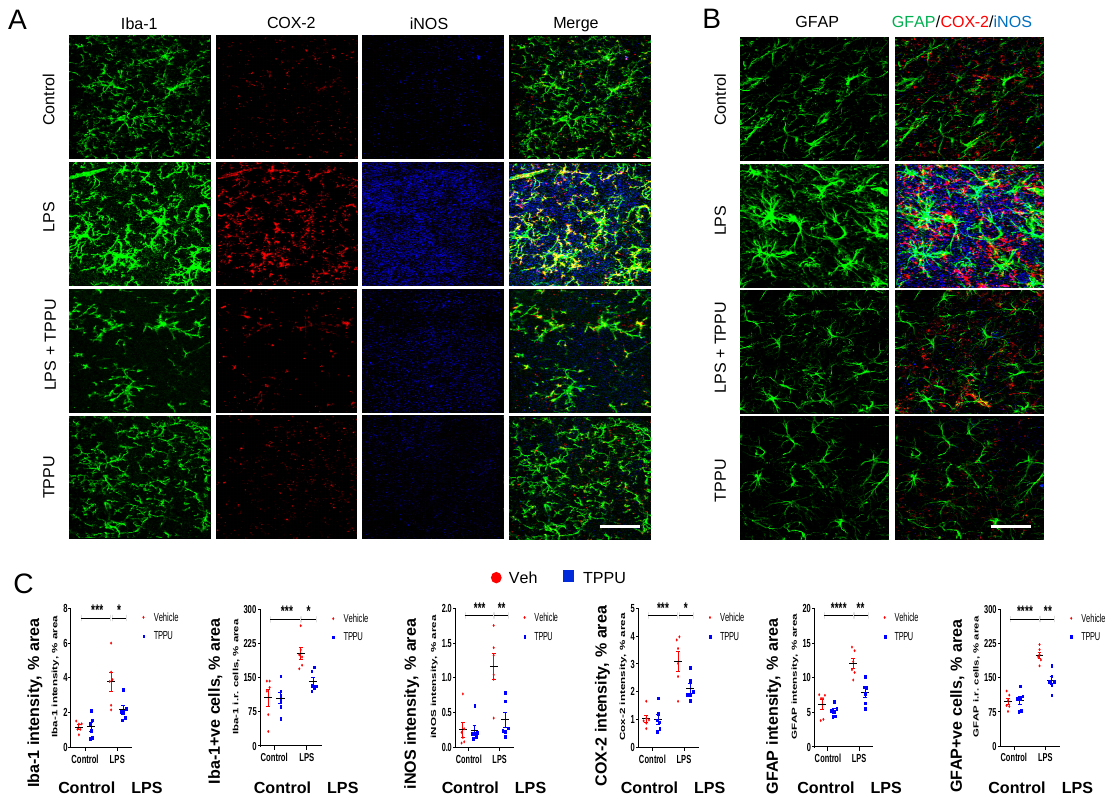

**Fig. S8. TPPU mitigates LPS-induced inflammation in C57BL/6 mice.** (A) Triple immunofluorescence staining of Iba-1 (green), COX-2 (red) and iNOS (blue) in hippocampus of mice treated with LPS, TPPU or LPS + TPPU as outlined in Fig. S7a. Scale bar, 100 μm. (B) Immunofluorescence staining of GFAP (left) and merged panel of GFAP (green), COX-2 (red) and iNOS (blue) (right). Scale bar, 75 μm. (C) Quantification of Iba-1, iNOS, COX-2 and GFAP intensities and Iba-1+ve and GFAP+ve cells in the hippocampus. Data are means ± SEM of six to eight mice per group. ****P < 0.0001, ***P < 0.001, **P < 0.01, *P < 0.05. Data were analyzed by one-way ANOVA with Tukey’s multiple comparison test.

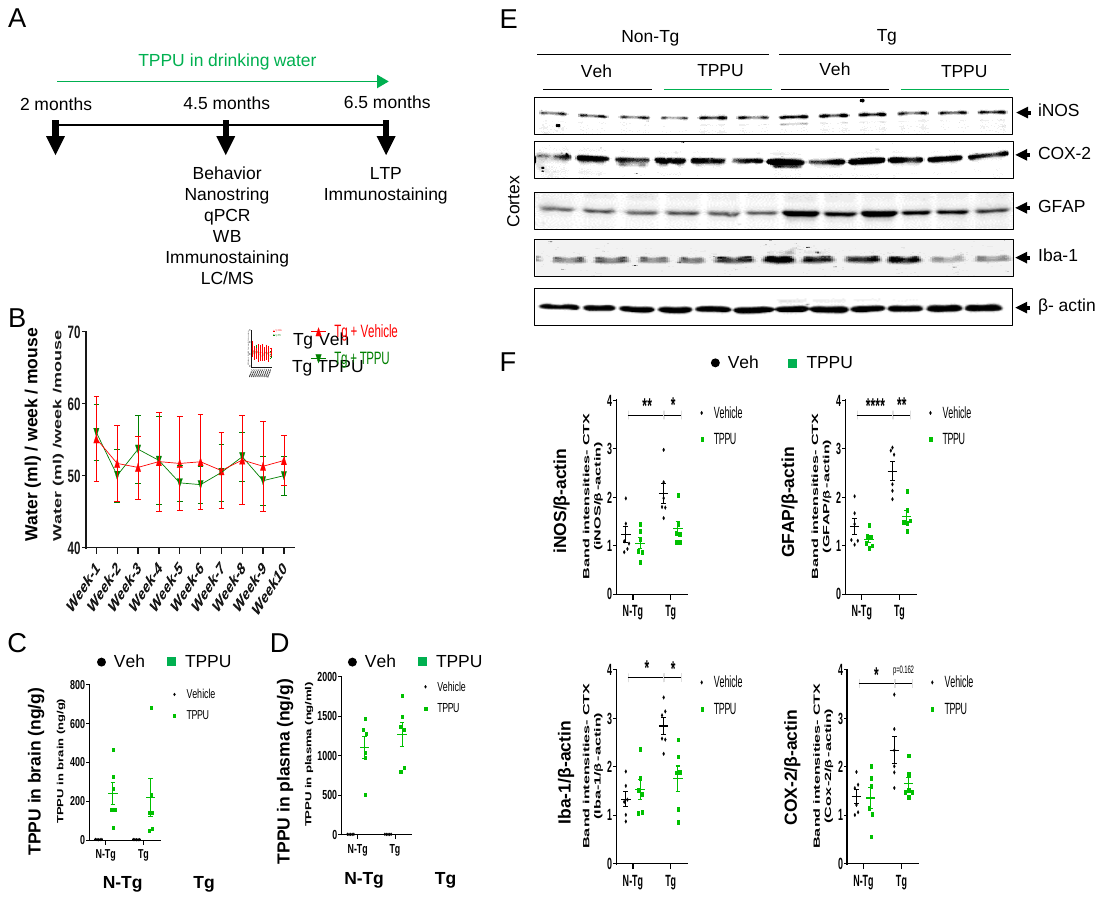

**Fig. S9.** **TPPU reduces neuroinflammation in Tg mice.** (A) Schematic diagram showing TPPU treatment and analysis. TPPU treatment starts at 2 months of age and continues for either 2.5 months or 4.5 months. At 4.5 months of age mice underwent for behavioral experiments. Brains were processed for analyses as indicated. At 6.5 months of age mice were euthanized for LTP or immunostaining. (B) Weekly average drinking water consumption per Tg mouse giving either vehicle (Veh) or TPPU. (C) The brain homogenates or (D) plasma were extracted and were analyzed by LC/MS for concentrations of TPPU. (E) Representative Western blot of iNOS, COX-2, GFAP and Iba-1 in cortex. (F) Scattered dot plot showing mean Western blot protein expression over β-actin. Data are means ± SEM of six to eight mice per group. ****P < 0.0001, **P < 0.01, *P < 0.05. Data were analyzed by two-way ANOVA with Bonferroni’s multiple comparison test.

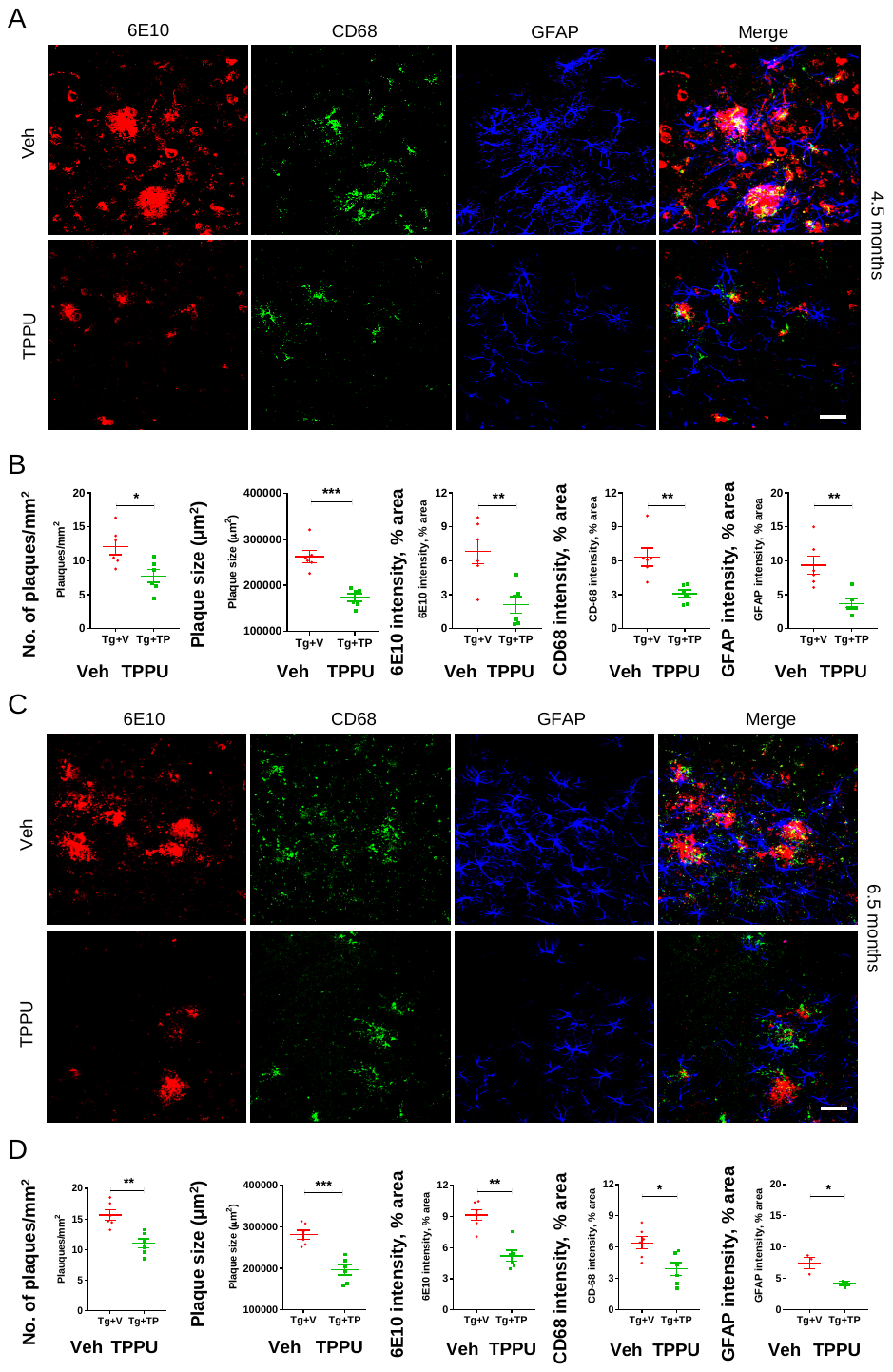

**Fig. S10. TPPU reduces Aβ burden in Tg mice.** (A) Triple immunofluorescence staining of 6E10 (red), CD68 (green) and GFAP (blue) in the cortex of 4.5-month-old mice (2 months treatment). (B) Quantification of 6E10+ve plaque number, size and 6E10 and CD68 intensities. (C) Triple immunofluorescence staining of 6E10 (red), CD68 (green) and GFAP (blue) in the cortex of 6.5-month-old mice (4.5 months treatment). (D) Quantification of 6E10+ve plaque number, size and 6E10 and CD68 intensities. Scale bars, 200 μm. Data are means ± SEM of six mice per group. ***P < 0.001, **P < 0.01, *P < 0.05. Data were analyzed by Student’s *t*-test.

**Table S1. P-glycoprotein (P-gp) substrate evaluation of TPPU in Caco-2 cell line**

| **Compound ID** | **Concentration (μM)** | **verapamil (μM)** | **P_app (A-B)_** | **P_app (B-A)_** | **Efflux Ratio** | **Recovery (%)** | **Recovery (%)** |
| --- | --- | --- | --- | --- | --- | --- | --- |
|  |  |  | **(10^-6^, cm/s)** | **(10^-6^, cm/s)** |  | **AP-BL** | **BL-AP** |
| Propranolol | 5 | - | 25.31 | 15.94 | 0.63 | 74.93 | 85.28 |
|  | 5 | 100 | 30.78 | 19.84 | 0.64 | 89.34 | 94.12 |
| Digoxin | 5 | - | 0.35 | 19.70 | 56.24 | 94.90 | 94.54 |
|  | 5 | 100 | 4.52 | 6.06 | 1.34 | 109.30 | 98.99 |
| TPPU | 5 | - | 18.03 | 24.45 | 1.36 | 80.88 | 85.49 |
|  | 5 | 100 | 19.50 | 19.29 | 0.99 | 80.75 | 84.75 |

**Table S2. Human demographics related to “Human subjects” in methods**

| **INDDID** | **Diagnosis** | **Sex** | **Age** |
| --- | --- | --- | --- |
| 109650 | AD | Female | 64 |
| 115165 | AD | Male | 64 |
| 120851 | AD | Male | 64 |
| 106029 | AD | Male | 71 |
| 121576 | AD | Male | 69 |
| 101034 | AD | Male | 67 |
| 116620 | AD | Female | 55 |
| 112723 | AD | Male | 72 |
| 117504 | Normal | Male | 59 |
| 113695 | Normal | Female | 59 |
| 102215 | Normal | Female | 65 |
| 100786 | Normal | Male | 61 |
| 101799 | Normal | Male | 70 |
| 103376 | Normal | Female | 68 |
| 107712 | Normal | Male | 70 |
| 110602 | Normal | Male | 55 |
| 113818 | Normal | Female | 65 |
| 116519 | Normal | Female | 67 |

| **Gene** | **Forward (5’-3’)** | **Reverse (3’-5’)** |
| --- | --- | --- |
| mEphx2 | CCACTCTAGGCCACAGCCTT | CCAAGCAGGAAGTCTCTGGAAA |
| hEphx2 | GTGACCGGAATCCAGCTTCTCAATA | CCAAGAATACCAACTCTCGGGAAAT |
| mCyp2c39 | GAGGAAGCATTCCAATGGTAGAA | TGTGAAGCGCCTAATCTCTTTC |
| mCyp2j5 | TCTGGGAAGCACTCCATCTCA | CCCTGGTGGGTAGTTTTTGG |
| mCyp2j9 | TGGCTGATTTCCTCAAAAACCG | ACTGCTGAAGGGATAGGTGGG |
| hFaah | CTCTGCTGCCAAGGCTGT | TGCAGTTCCCAGAGTTTTCC |
| hCyp4f81 | CATCTTCAGCTTTGACAGCAA | TGAGCTCCATGATCGCAGTA |
| hPla2g2a | ACCTGCCCTGTCTCCAAAC | TTTGTTCTGCACTCCTGCTC |
| hPlag7 | TGGCTCTACCTTAGAACCCTGA | TTTTGCTCTTTGCCGTACCT |
| mIL-1α | CGCTTGAGTCGGCAAAGAAAT | CTTCCCGTTGCTTGACGTTG |
| mIL-1β | GCAACTGTTCCTGAACTCAACT | ATCTTTTGGGGTCCGTCAACT |
| mIL-6 | TAGTCCTTCCTACCCCAATTTCC | TTGGTCCTTAGCCACTCCTTC |
| mTNF-α | CCCTCACACTCAGATCATCTTCT | GCTACGACGTGGGCTACAG |
| miNOS | CCCTTCCGAAGTTTCTGGCAGCAGC | GGCTGTCAGAGCCTCGTGGCTTTGG |
| mC3 | AAG CAT CAA CAC ACC CAA CA | CTT GAG CTC CAT TCG TGA CA |
| mGfap | AGAAAGGTTGAATCGCTGGA | CGGCGATAGTCGTTA |
| mAif-1 | CAGACTGCCAGCCTAAGACA | AGGAATTGCTTGTTGATCCC |
| mCcl-2 | GAAGGAATGGGTCCAGACAT | ACGGGTCAACTTCACATTCA |
| mCcl-3 | CTCCCAGCCAGGTGTCATTTTC | AGGCATTCAGTTCAGGTCAG |
| mCcl-4 | CCAACTTCCTGCTGTTTCTCT | GTCTGCCTCTTTTGGTCAGGA |
| mCcl-5 | GTGCCCACGTCAAGGAGTAT | CCACTTCTTCTCTGGGTTGG |
| mCxcl-1 | ACCCAAACCGAAGTCATAGCC | TTGTCAGAAGCCAGCGTT |
| mCxcl-10 | CCCACGTGTTGAGATCATTG | CACTGGGTAAAGGGGAGTGA |
| mGbp-2 | GGGGTCACTGTCTGACCACT | GGGAAACCTGGGATGAGATT |
| mAspg | GCTGCTGGCCATTTACACTG | GTGGGCCTGTGCATACTCTT |
| mSerping | ACAGCCCCCTCTGAATTCTT | GGATGCTCTCCAAGTTGCTC |
| mAmigo-2 | GAGGCGACCATAATGTCGTT | GCATCCAACAGTCCGATTCT |
| m18s | CCATTCGAACGTCTGCCCTAT | GTCACCCGTGGTCACCATG |
| hGapdh | AGGGCTGCTTTTAACTCTGGT | CCCCACTTGATTTTGGAGGGA |
| mCD68 | ACTGGTGTAGCCTAGCTGGT | CCTTGGGCTATAAGCGGTCC |
| mCox-2 | TTCAACACACTCTATCACTGGC | AGAAGCGTTTGCGGTACTCAT |
| mC1q | AAAGGCAATCCAGGCAATATCA | TGGTTCTGGTATGGACTCTCC |
| hCyp2j2 | ACTGTCGCCTTTCTGCTCG | CTGCTCGAAGTCCACAAGGAA |
| hCyp2c8 | AGATCAGAATTTTCTCACCC | AACTTCGTGTAAGAGCAACA |
| hCyp2c19 | TGCTCTCCTTCTCCTGCTGAAG | TGCCAACGACACGTTCAATC |
| hCox-2 | GAATGGGGTGATGAGCAGTT | CAGAAGGGCAGGATACAGC |

**Table S3. Primer sequences for qPCR. Related to “RNA extraction and quantitative PCR (qPCR)”. Prefix ‘h’ denotes human and ‘m’ denotes mouse.**
